# Claustral pathway coordinates distributed cortical dynamics with hippocampal output during sleep to promote memory consolidation

**DOI:** 10.64898/2026.08.13.744575

**Authors:** Coline Portet, Flora Thellier, Matthieu Aguilera, Thomas Blondel, Karin Herbeaux, Caroline Mursch, Jesse Jackson, Demian Battaglia, Yaroslav Sych, Romain Goutagny

**Author notes:** CP and FT are co-first authors who contributed equally to this work. Their order of appearance is alphabetical and does not reflect relative contribution; either author may therefore cite this article with their name listed first in bibliographic references. **Correspondence and requests for materials** should be addressed to Romain Goutagny.

## Abstract

The claustrum is a broadly connected subcortical structure proposed to coordinate distributed cortical activity. Claustral neurons are recruited during synchronized brain states and contribute to sleep-dependent memory consolidation, primarily through effects on cortical dynamics. Yet the claustrum also innervates the subicular complex, a major hippocampal output node, raising the possibility that it may regulate both cortical state and hippocampo-cortical dialogue during sleep. Here, we identify a projection-defined population of claustral neurons targeting the subicular complex, CLAsc, that is preferentially recruited during slow-wave sleep and tracks cortical and subicular sleep dynamics. Optogenetic activation of CLAsc neurons during post-learning sleep enhanced spatial memory consolidation without detectable changes in ripple occurrence or slow-oscillation-spindle coupling. Instead, CLAsc activation imposed a stereotyped cortical-subicular motif: a brief gamma-rich Up state followed by a coordinated Down state across prefrontal, retrosplenial and subicular regions. Within this gamma-rich Up state, interareal coherence increased, gamma bursts became synchronized, and lagged directed interactions were transiently reorganized, including an enhanced prefrontal-to-subicular component. Together, these findings identify the claustrum as a state-dependent coordinator that transforms ongoing cortical and hippocampal-output activity into coordinated network transitions during sleep, providing a circuit mechanism through which distributed brain states may support memory consolidation.

## Introduction

The claustrum is a thin subcortical structure best known for its exceptionally dense and widespread reciprocal connectivity with the cerebral cortex ^1^. This architecture has placed the claustrum at the center of theories proposing that it coordinates distributed cortical activity, including models of sensory integration, attention, salience processing, and cognitive control ^2–4^. More recently, this coordinating role has been linked to brain-state regulation. Claustral neurons are preferentially recruited during rest, slow-wave sleep, and other states marked by strong cortical synchrony, and claustral activation can suppress ongoing cortical activity through recruitment of cortical inhibitory interneurons ^5–11^. These findings suggest that claustro-cortical signaling may help organize activity during synchronized brain states. Consistent with this idea, a retrosplenial-projecting claustral population was recently shown to shape cortical slow-wave dynamics and facilitate memory consolidation during slow-wave sleep ^9^. Sleep-dependent memory consolidation, however, relies on the coordinated exchange of information between the hippocampal formation and distributed cortical networks ^12,13^. How claustral modulation of cortical sleep dynamics contributes to this dialogue remains unclear, particularly because the claustrum does not project to the hippocampus proper ^14^. A potential route has nevertheless been largely overlooked. Projection-defined and brain-wide tracing studies have shown that claustral neurons innervate the subicular complex, a principal output structure of the hippocampal formation, while collateralizing across broad cortical territories ^14–17^. This anatomical organization places the claustrum in a unique position to coordinate distributed cortical activity with hippocampal output during sleep. Whether this pathway is recruited during slow-wave sleep and contributes to memory consolidation remains unknown.

Here, we tested this possibility by targeting subicular-projecting claustral neurons, which we refer to as CLAsc. Combining projection-specific viral targeting, brain-wide anatomical mapping, fiber photometry, multisite electrophysiology, and sleep-restricted optogenetic manipulation in freely moving mice, we show that CLAsc neurons are preferentially recruited during slow-wave sleep, track cortical and subicular synchronization across multiple timescales, and promote memory consolidation when activated during post-learning sleep. CLAsc activation evokes a stereotyped network motif consisting of a brief distributed gamma-rich Up state followed by a coordinated Down state spanning prefrontal, retrosplenial, and subicular regions. Within the gamma rich Up state, interareal coherence increases and directed interactions are transiently reorganized along a cortex-to-subiculum axis. These findings identify a subicular-projecting claustral pathway that coordinates cortical dynamics with hippocampal output during slow-wave sleep and supports sleep-dependent memory consolidation.

## Results

### Subicular-projecting claustral neurons are positioned to coordinate hippocampo-cortical dynamics during sleep

Claustral neuronal populations are organized by projection target, and recent work has shown that a retrosplenial-projecting subpopulation shapes cortical slow-wave dynamics and facilitates memory consolidation during sleep ^9^. Here we focused on neurons projecting to the subicular complex (CLAsc), a population anatomically positioned at the interface between hippocampal output and distributed cortical territories.

To selectively label this population, we combined a retrograde virus expressing Cre recombinase injected into the subicular complex with a Cre-dependent eYFP virus injected into the claustrum (Fig. 1A). Immunostaining for TLE4, a transcription factor expressed in adjacent cortical areas but absent from claustral neurons ^18^, confirmed that eYFP-positive cells were confined to the claustral core and did not overlap with TLE4+ cells (Fig. 1A), validating the specificity of the intersectional labeling strategy.

**Figure 1.**
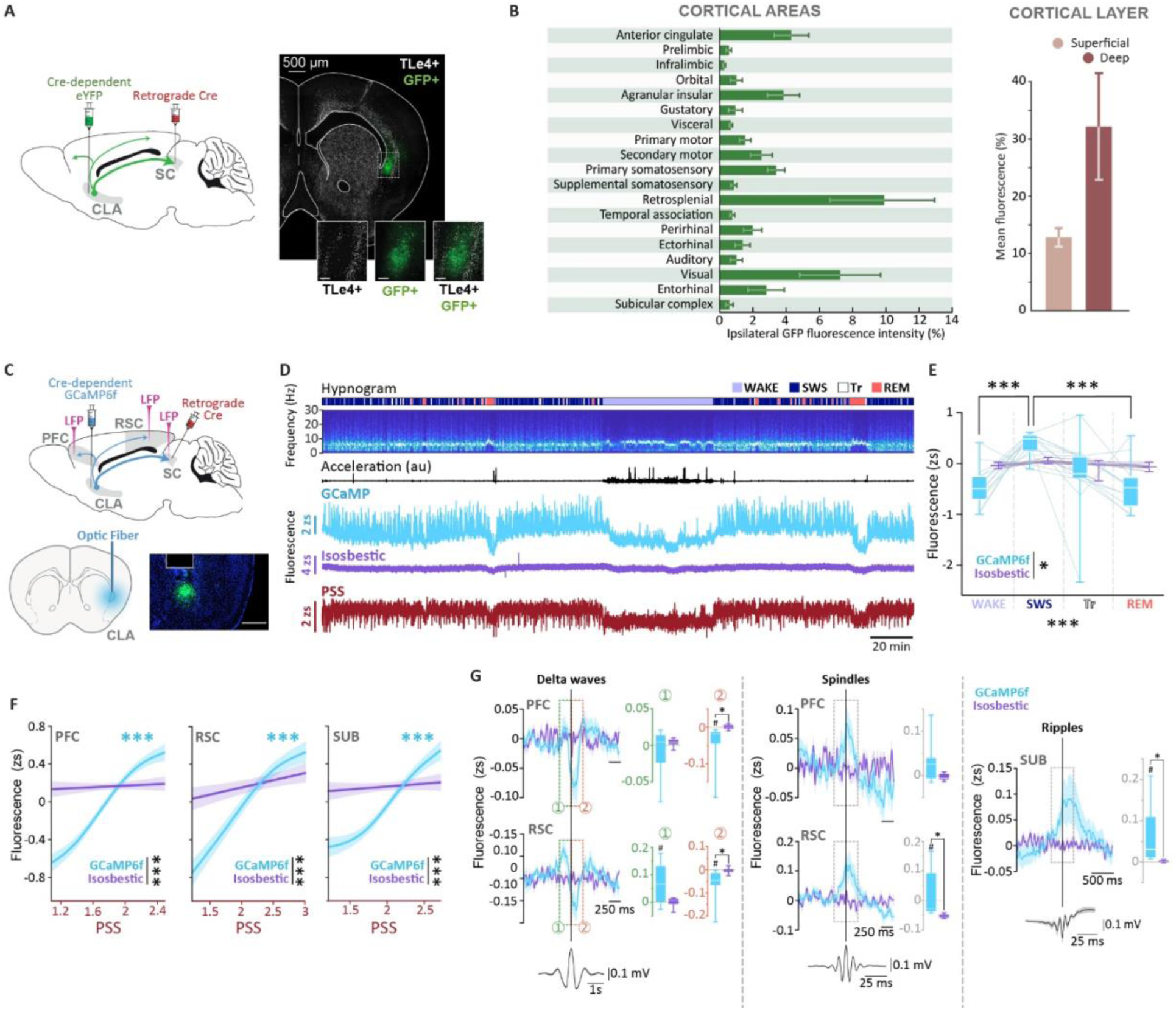
Anatomical and functional characterization of subicular-projecting claustral neurons (CLAsc). **A.** Viral strategy and representative sections showing retrograde-Cre-dependent eYFP expression in subicular-projecting claustral neurons. GFP-positive neurons were confined to the claustral core and excluded from adjacent TLE4-positive cortex. Scale bars: 500 µm (whole hemisphere), 125 µm (claustrum insert). **B.** Ipsilateral distribution of CLAsc axonal fluorescence across cortical and subicular territories. Inset, cortical fluorescence was enriched in deep layers. Data are mean ± SEM, n = 3 mice. **C.** Configuration for CLAsc fiber photometry and simultaneous LFP recordings from PFC, RSC, and SUB. Scale bars: 400 µm. **D.** Representative recording showing vigilance state, PFC spectrogram, movement, CLAsc GCaMP6f signal, isosbestic control, and PFC power-spectrum slope. **E.** CLAsc GCaMP6f fluorescence was increased during SWS relative to Wake and REM, whereas the isosbestic signal was not state modulated. Signal × state interaction: F(2,48.2) = 5.31, p = 0.008; n = 7 mice. **F.** CLAsc GCaMP6f fluorescence increased with local power-spectrum slope in PFC, RSC, and SUB during SWS, whereas the isosbestic signal showed little modulation. Lines show GAMM fits and shaded areas indicate SEM. all p < 0.0001. **G.** CLAsc fluorescence aligned to cortical delta waves and spindles and to subicular ripples during SWS. Traces show mean ± SEM GCaMP6f and isosbestic signals aligned to event center; grey boxes indicate the predefined quantification windows. GCaMP6f fluorescence differed from the isosbestic signal following RSC delta waves and around subicular ripples, and was greater than zero before RSC delta waves and around RSC spindles. Representative event waveforms are shown below. Paired and one-sample Wilcoxon signed-rank tests; n = 7 for delta waves and spindles, n = 6 for ripples. *p < 0.05, GCaMP6f versus isosbestic; #p < 0.05, GCaMP6f versus zero.

Quantification of fluorescence density across cortical and subcortical regions using the Automated Brain-wide Alignment and Annotation pipeline (ABBA ^19^; n = 3 mice) revealed that CLAsc projections are distributed broadly across the cortical mantle, with prominent outputs to cortical territories including prefrontal, insular, retrosplenial, visual, and entorhinal cortices, as well as to the subicular complex (Fig. 1B). CLAsc projections preferentially innervated deep cortical layers (L5-L6) over superficial layers (L1-L4; Fig. 1B, inset). This laminar bias positions CLAsc outputs within layers enriched in long-range projection neurons, consistent with a role in interareal and subcortical communication. Together, these anatomical features identify CLAsc neurons as a candidate pathway through which claustral activity could interface with distributed cortical and hippocampal-output networks.

We next examined how CLAsc activity varies across vigilance states. Using the same dual-viral strategy, we expressed the calcium indicator GCaMP6f in CLAsc neurons and recorded fluorescence through an optic fiber implanted above the claustrum together with local field potentials (LFPs) from the prefrontal (PFC) and retrosplenial (RSC) cortices and the subicular complex (SUB) (n = 7 mice; Fig. 1C). Recordings in freely moving mice revealed that CLAsc calcium activity was significantly higher during SWS than during wakefulness or REM sleep (linear mixed-effects model, main effect of state: p < 0.001; Post hoc comparisons within the GCaMP signal: SWS > wake, p < 0.001; SWS > REM, p < 0.001; n = 7 mice; Fig. 1D, E). This modulation was absent in the isosbestic control signal, indicating that it did not reflect movement- or hemodynamics-related artifacts.

To relate CLAsc activity to cortical synchronization, we computed the power-spectrum slope (PSS), an index of the relative contribution of slow versus fast frequencies. Generalized additive mixed models (GAMMs ^20^) revealed a significant positive relationship between GCaMP fluorescence and PSS across all recorded regions (PFC: p <2.2 ×10⁻^16^, edf = 3.56; RSC: p = 2.2 ×10⁻^16^, edf = 2.83; SUB: p < 2.2 ×10⁻¹⁶, edf = 3.47; Fig. 1F), indicating that CLAsc neurons are most active during synchronized, delta-dominated states. No significant relationship was observed for the isosbestic signal in any region (all p > 0.26), and direct comparison of GCaMP and isosbestic smooths confirmed that their PSS dependence differed significantly across all regions (GAMMs; PFC: p = 2.2 ×10⁻^16^; RSC: p = 1.993 ×10⁻^15^; SUB: p < 2.2 ×10⁻^16^). Similar results were obtained using the theta/delta ratio as an alternative index of cortical synchronization (Fig. S1). We next asked whether CLAsc activity was temporally organized around the discrete oscillatory events of SWS. Event-triggered analyses revealed the clearest modulation around subicular ripples and retrosplenial delta waves (Fig. 1G). Around subicular ripples, GCaMP fluorescence increased within the predefined -250 to +250 ms window and differed from the isosbestic signal (paired Wilcoxon signed-rank test, p = 0.031; GCaMP versus zero, p = 0.031; isosbestic versus zero, p = 0.219; n = 6). Following RSC delta waves, GCaMP fluorescence decreased during the subsequent 250 ms, both relative to the isosbestic signal and to zero (both p = 0.016; n = 7). GCaMP fluorescence was also greater than zero before RSC delta waves (p = 0.047) and around RSC spindles (p = 0.016), although neither effect differed significantly from the isosbestic control (p = 0.078 and p = 0.156, respectively). No significant modulation was detected around PFC spindles or delta waves.

Together, these results indicate that CLAsc recruitment tracks both the ongoing level of cortical synchronization and the occurrence of discrete SWS events, including cortical delta waves and spindles and subicular ripples.

### CLAsc activation during post-learning SWS enhances spatial memory consolidation

Having established that CLAsc neurons are preferentially recruited during synchronized sleep states, we next asked whether activating this pathway during post-learning sleep influences memory consolidation. We expressed the excitatory opsin ChETA bilaterally in CLAsc neurons and implanted optical fibers above the claustrum, together with LFP electrodes in PFC, RSC and SUB for online sleep monitoring (n = 10 mice; Fig. 2A). Mice performed an object-location task in which the sampling period was deliberately kept short (4 min), so that memory performance after 24 h was near chance, thereby increasing sensitivity to manipulations that facilitate consolidation (Fig. 2B). During the first hour of post-learning sleep, 5 ms light pulses were delivered at 0.5 Hz during online-detected SWS. In control sessions, the same animals received red light with identical timing. Offline scoring confirmed that stimulation was effectively restricted to SWS, with 89 ±5% of light pulses occurring during verified SWS epochs and no mouse receiving less than 78% of stimulations during SWS (Fig. 2B).

**Figure 2.**
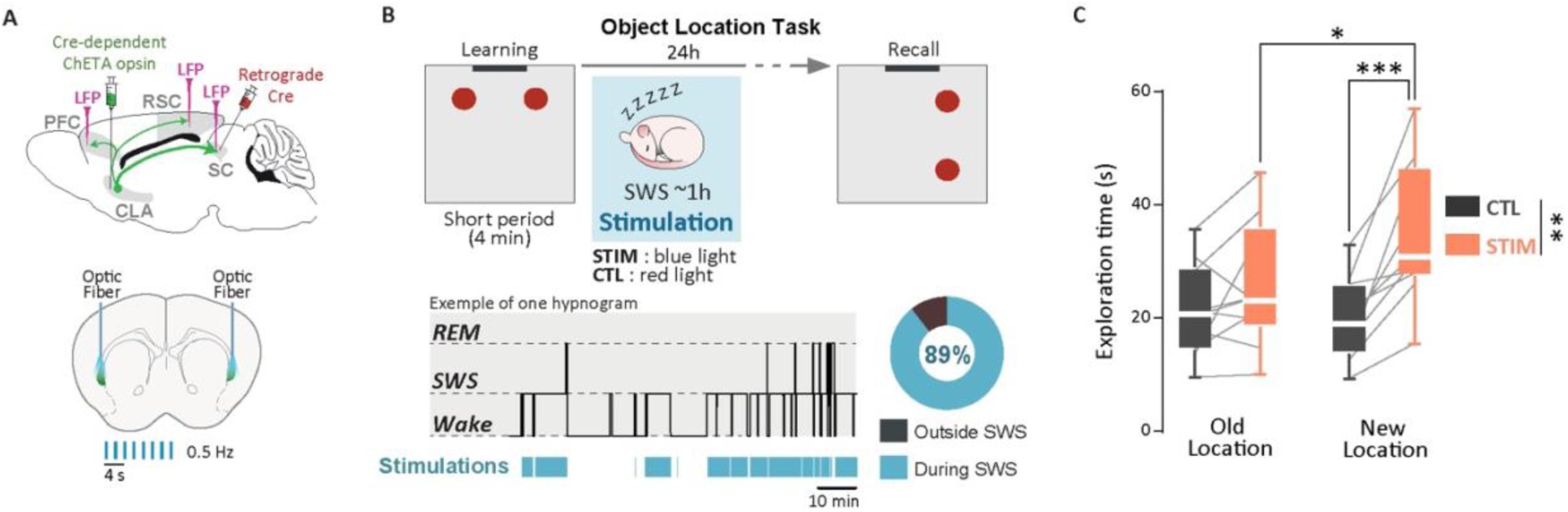
CLAsc activation during post-learning sleep enhances spatial memory consolidation. **A.** Optogenetic strategy for activating CLAsc neurons while recording LFPs from PFC, RSC, and SUB. Blue-light stimulation was delivered as 5-ms pulses at 0.5 Hz; red-light sessions served as controls. **B.** Object-location task and sleep-stimulation protocol. Mice explored two identical objects for 4 min, received CLAsc stimulation during the first hour of post-learning SWS, and were tested 24 h later with one object displaced. Representative hypnogram shows stimulation timing. Across mice, 89 ±5% of stimulations occurred during verified SWS. **C.** Recall-phase object exploration. CLAsc stimulation selectively increased exploration of the displaced object. Object × Condition interaction: F_(1,18)_ = 9.30, p = 0.007; displaced versus non-displaced object in STIM, p = 0.009; CTL, p = 0.918. Lines indicate paired mice. Data are min-to-max box plots with individual values overlaid. *p < 0.05, **p < 0.01.

CLAsc activation improved memory performance at recall (Fig. 2C). A repeated-measures ANOVA on test-phase exploration time revealed a significant main effect of Object (F_(1,18)_ = 4.60, p = 0.046, ηp² = 0.203) and a significant Object × Condition interaction (F_(1,18)_ = 9.30, p = 0.007, ηp² = 0.341). Post-hoc comparisons showed that mice preferentially explored the displaced object in the STIM condition (p = 0.009), but not in the CTL condition (p = 0.918). Exploration of the displaced object was also higher in STIM than in CTL sessions (p = 0.023), whereas exploration of the non-displaced object did not differ between conditions (p = 1.000). Consistent with this effect, the memory index was significantly higher in the STIM than in the CTL condition (paired t_(9)_ = 2.55, p = 0.031, Cohen’s d = 0.8), and differed from chance only in the STIM condition (one-sample t_(9)_ = 3.87, p = 0.002), not in the CTL condition (one-sample t_(9)_ = 0.66, p = 0.74).

Total object exploration during the sampling phase was unchanged between conditions (paired t_(9)_ = 0.540, p = 0.54, Cohen’s d = 0.2), arguing against differences in initial exploration. CLAsc stimulation also did not alter global sleep architecture or bout structure during the first two post-learning hours, making sleep disruption an unlikely explanation for the behavioral effect (Fig. S2). Together, these results demonstrate that selective activation of CLAsc neurons during post-learning SWS is sufficient to enhance spatial memory consolidation.

### CLAsc activation evokes a stereotyped gamma-rich Up state followed by a Down state

To characterize the network dynamics underlying CLAsc-dependent memory consolidation, we examined stimulation-evoked responses during SWS using simultaneous recordings from PFC, RSC and SUB (n = 8 mice). Blue-light activation of CLAsc neurons evoked a highly stereotyped response across all three regions, visible in the averaged LFP as a prominent slow deflection (Fig. 3A). Time-frequency analysis revealed that this slow response was preceded by a brief increase in gamma-band power, peaking within the first tens of milliseconds after stimulation, followed by a suppression of high-frequency activity and a slower component in the delta range. The MUA envelope decreased after the gamma burst, consistent with the emergence of a population Down state. This biphasic gamma-to-slow-wave response was absent or markedly weaker in red-light control sessions. Cluster-based permutation tests confirmed significant stimulation-evoked LFP responses in all regions, with significant clusters in PFC (p = 0.03 and p = 0.0075), RSC (p = 0.03, p = 0.01, p = 0.04 and p = 0.01), and SUB (p = 0.0075, p = 0.04, p = 0.04 and p = 0.03; Table S1). The Kuramoto index revealed a sharp transient increase in phase consistency across stimulations in all three regions (PFC: p = 0.0075; RSC: p = 0.0075; SUB: p = 0.0075 and p = 0.047; Fig. 3A, Table S1), confirming that CLAsc activation generates a reproducible, time-locked network response rather than a variable or region-specific effect.

**Figure 3.**
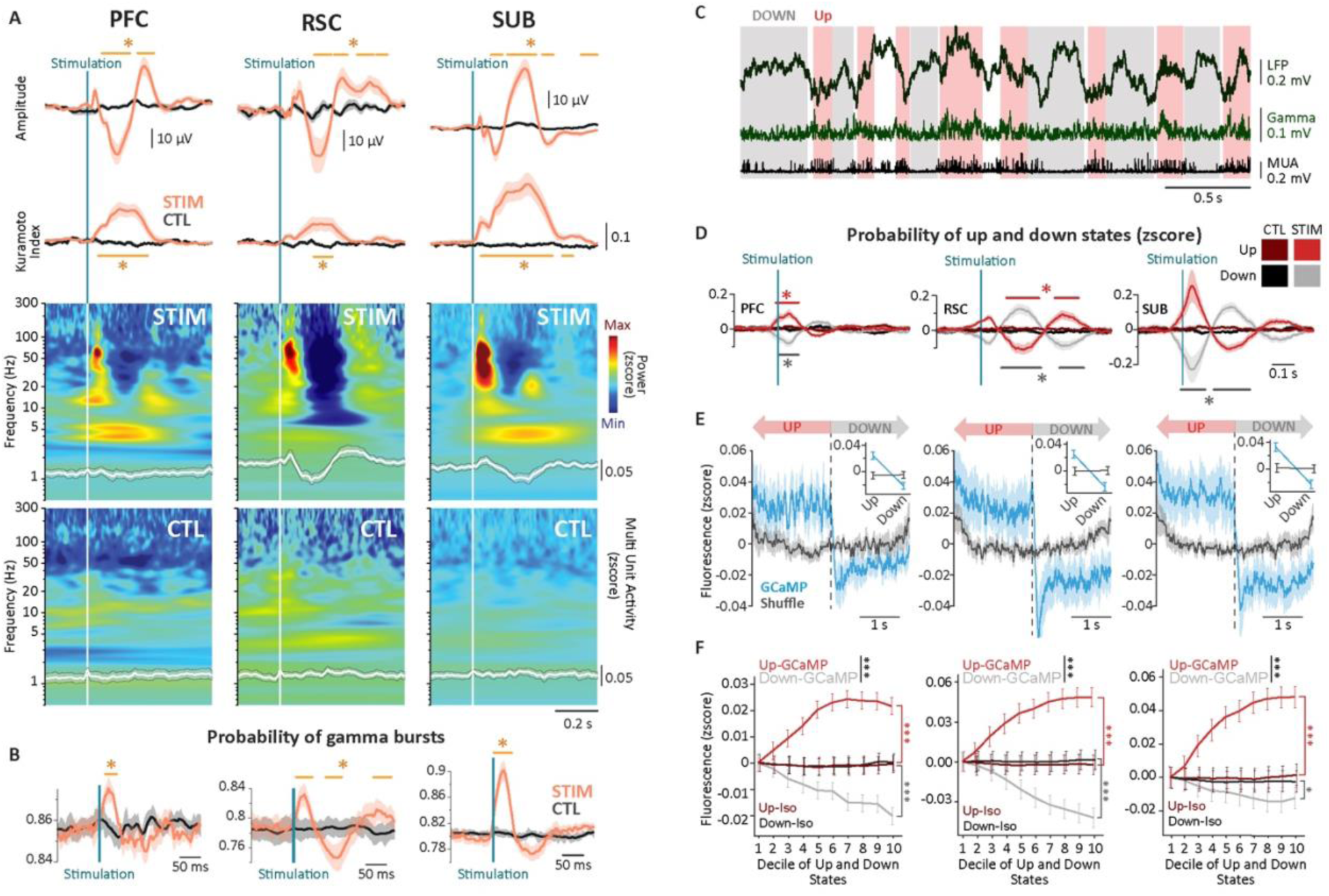
CLAsc activation evokes a gamma-rich Up state followed by a Down state. **A.** Stimulation-evoked responses in PFC, RSC, and SUB during SWS. Top, average LFP traces and Kuramoto index aligned to light onset for CLAsc stimulation and control trials. Bottom, stimulation-aligned time-frequency maps with MUA overlaid in white. Orange bars indicate significant STIM versus CTL clusters. **B.** Gamma-burst probability aligned to stimulation onset. CLAsc activation transiently increased gamma-burst probability in PFC, RSC, and SUB, followed by a brief reduction. Shaded areas indicate SEM; orange bars indicate significant STIM versus CTL clusters. **C.** Example of Up and Down state detection from LFP, gamma-band activity, and MUA. Up and Down states are indicated by red and grey shading, respectively. **D.** Stimulation-aligned Up and Down state probabilities in PFC, RSC, and SUB. CLAsc activation increased Up-state probability shortly after stimulation and was followed by a delayed increase in Down state probability. Shaded areas indicate SEM; bars indicate significant STIM versus CTL clusters. **E.** CLAsc calcium activity aligned to spontaneous Up-to-Down transitions detected in PFC, RSC, and SUB during SWS. GCaMP6f fluorescence decreased at the Up-to-Down transition, whereas shuffled controls showed no comparable modulation. Insets show mean fluorescence during Up and Down states. **F.** CLAsc fluorescence across normalized Up and Down states. GCaMP6f activity increased across Up states and decreased across Down states, whereas the isosbestic signal remained weakly modulated. Data are mean ± SEM across state deciles. Significance levels are indicated in the panels; *p < 0.05, **p < 0.01, ***p < 0.001.

We next asked whether the early increase in gamma-band power reflected discrete transient events rather than a nonspecific broadband spectral change. To isolate gamma activity from the broadband LFP, we decomposed the signal using ensemble empirical mode decomposition (EEMD ^21,22^; Fig. S3) and reconstructed a gamma composite by combining the intrinsic mode functions spanning the gamma-frequency range. We then quantified the time-resolved probability of gamma-burst occurrence ^23^ in this EEMD-derived signal relative to stimulation onset (Fig. 3B, Fig. S4). CLAsc activation induced a short-latency increase in gamma-burst probability in all recorded regions, followed by a transient suppression. Cluster-based permutation tests confirmed significant post-stimulation increases in gamma-burst occurrence in PFC (p = 0.0075), RSC (p = 0.02, p = 0.03 and p = 0.04), and SUB (p = 0.0075; Table S1). Thus, the early response to CLAsc activation corresponds to a discrete gamma transient that precedes the subsequent reduction in population activity. Importantly, CLAsc activation did not increase cortical spindle probability or subicular ripple probability relative to stimulation onset, although it was followed by a delayed transient increase in delta-wave probability in SUB, with no corresponding change in PFC or RSC (Fig. S5). These results indicate that the stimulation-evoked gamma transient is not explained by an increased occurrence of canonical cortical spindles or subicular ripples.

Because this sequence suggested a transition from active to silent network states, we next quantified Up and Down states from multi-unit activity using a probabilistic state-space model ^24^ (Fig. S6) and asked how their probabilities were affected by stimulation (Fig. 3C). CLAsc activation transiently increased Up-state probability shortly after stimulation, followed by a delayed increase in Down-state probability approximately 200 ms later. For Up-state probability, cluster-based permutation tests revealed significant stimulation effects in PFC (p = 0.0075) and RSC (p = 0.037 and p = 0.01; Fig. 3D and Table S1). For Down-state probability, significant clusters were detected in PFC (p = 0.0075), RSC (p = 0.01 and p = 0.04), and SUB (p = 0.0075 and p = 0.047; Fig. 3D; Fig. S7 and Table S1). This sequential Up-to-Down modulation mirrors the gamma-to-slow-wave response observed in the LFP and indicates that CLAsc activation biases ongoing SWS activity toward a structured state transition.

To test whether the evoked motif reflects a physiological mode of CLAsc operation, we aligned photometry signals to spontaneous Up-to-Down transitions during SWS (Fig. 3E). Event-aligned analyses further revealed a sharp decrease in spontaneous CLAsc activity at the transition into Down states in all three regions (Linear mixed model based on 2 seconds window; post-hoc comparison *GCaMP-Shuffle-Up vs. GCaMP-Shuffle-Down*: PFC p < 0.001, RSC p < 0.001, SUB p < 0.001, n = 7), whereas the isosbestic control signal did not show corresponding Up-Down modulation. When Up and Down episodes were normalized by their relative duration, GCaMP activity progressively increased across Up states and decreased across Down states, whereas the isosbestic signal remained flat (Fig. 3F). Thus, endogenous CLAsc activity is temporally organized across the Up-Down cycle, rising during active network states and falling rapidly as the network enters the Down state. Its modulation around spindles, ripples and delta waves further indicates that CLAsc activity is tightly coupled to ongoing cortical and subicular sleep dynamics.

Together, these results show that CLAsc activation drives a gamma-rich Up state followed by a coordinated Down state, and that this evoked sequence mirrors a physiological state-transition motif embedded in spontaneous CLAsc dynamics during SWS.

### CLAsc activation transiently coordinates hippocampo-cortical interactions during SWS

Having established that CLAsc activation evokes a gamma-rich Up state followed by a Down state in individual regions, we asked whether this motif was coordinated across the cortical-subicular network. CLAsc activation increased the probability of a synchronous Up state shortly after stimulation, followed by a delayed increase in synchronous Down-state probability across PFC, RSC and SUB (synchronous Up states: cluster p = 0.04 and p = 0.01; synchronous Down states: cluster p = 0.014 and p = 0.046; Fig. 4A, Fig. S7 and Table S1). Thus, the Up-to-Down sequence evoked by CLAsc activation emerges as a coordinated network motif spanning cortical and subicular sites. We then examined whether this coordinated motif corresponds to a physiological mode of CLAsc recruitment during spontaneous SWS. CLAsc photometry signals were aligned to naturally occurring Up and Down states detected simultaneously in PFC, RSC and SUB. Endogenous CLAsc activity was higher during synchronous Up states than during synchronous Down states, whereas the isosbestic control signal showed no comparable modulation (two-ways repeated-measures ANOVA; Post-hoc comparison: GCaMP Up and Down Kolmogorov-Smirnov distribution vs. GCaMP Shuffled KS distribution, adjusted p = 0.0029; Fig. 4B). This Up versus Down separation was greater than expected after temporal shuffling, indicating that CLAsc activity is specifically coupled to coordinated cortical-subicular state fluctuations. These results show that endogenous CLAsc activity is elevated during synchronous cortico-subicular Up states in SWS, consistent with the motif revealed by optogenetic stimulation.

**Figure 4.**
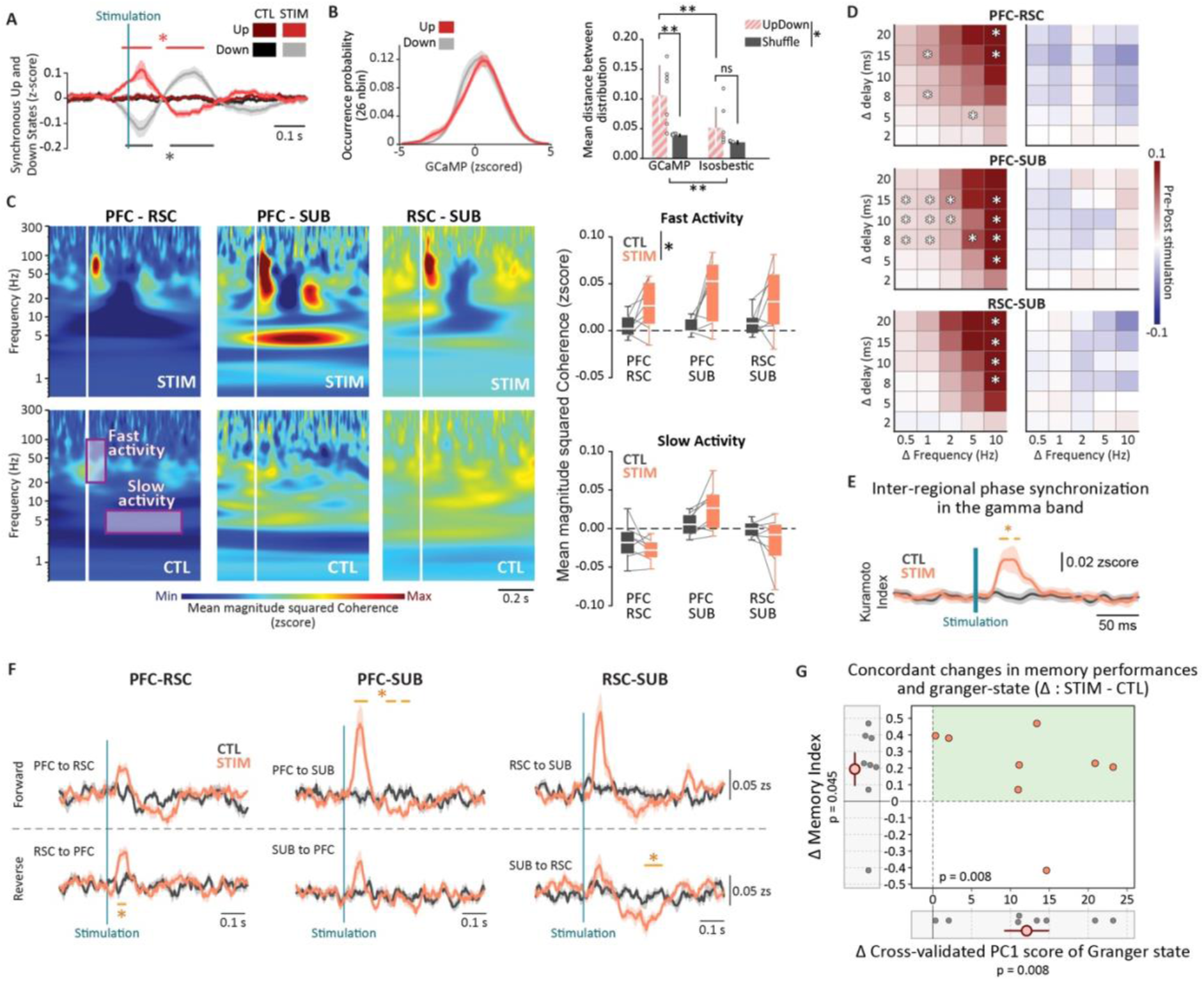
CLAsc activation transiently coordinates cortical-subicular interactions during SWS. **A.** Stimulation-aligned probability of synchronous Up and Down states across PFC, RSC, and SUB. Synchronous states were defined as periods during which the same state was detected simultaneously in all three regions. CLAsc activation increased synchronous Up-state probability shortly after stimulation, followed by a delayed increase in synchronous Down-state probability. Shaded areas indicate SEM; horizontal bars indicate significant STIM versus CTL clusters. **B.** Relationship between endogenous CLAsc activity and spontaneous synchronous Up and Down states. Left, distributions of GCaMP6f fluorescence during synchronous Up and Down states and their temporally shuffled controls. Right, Kolmogorov-Smirnov distance quantifying the separation between Up and Down-state fluorescence distributions. GCaMP6f showed greater Up-Down separation than shuffled controls, whereas the isosbestic signal did not. Bars show mean ± SEM with individual mice overlaid. **C.** Stimulation-aligned magnitude-squared wavelet coherence between PFC, RSC, and SUB. Heatmaps show coherence for each region pair in STIM and CTL sessions. Box plots quantify coherence within the early fast-frequency window (20-100 Hz, 10-100 ms) and the later slow-frequency window (3-7 Hz, 100-500 ms). Lines connect paired CTL and STIM values from the same mouse. **D.** Gamma-burst co-occurrence across region pairs. Heatmaps show the STIM-CTL difference in the change from pre- to post-stimulation co-occurrence as a function of temporal lag and frequency difference between bursts. Stars indicate delay-frequency bins surviving Benjamini-Hochberg false-discovery-rate correction. **E.** Cross-regional gamma-phase synchronization following CLAsc stimulation. The Kuramoto index was computed across PFC, RSC, and SUB from the instantaneous phase of the EEMD-derived gamma composite. CLAsc activation induced a transient increase in interregional gamma-phase alignment during the early post-stimulation window. Shaded areas indicate SEM; the horizontal bar indicates a significant STIM versus CTL cluster. **F.** Stimulation-aligned covariance-based Granger-causality analysis of gamma-band interactions between PFC, RSC and SUB, showing lagged directed interactions for each region pair and direction. CLAsc activation produced short-latency increases in lagged directed interactions, most prominently from RSC to PFC and from PFC to SUB. Shaded areas indicate SEM; horizontal bars indicate significant STIM versus CTL clusters. **G.** Relationship between stimulation-induced changes in the cross-validated Granger-causality network-state score and memory index. The network-state score was defined from the first principal component of the six directed Granger-causality measures using a leave-one-mouse-out procedure. Both measures are expressed as within-animal STIM-minus-CTL differences. Seven of eight mice showed concordant increases in both measures. An exact within-mouse permutation test confirmed a significant joint positive effect of stimulation (p = 0.008). The magnitudes of the physiological and behavioral changes were not significantly correlated across animals (Spearman’s ρ = -0.38, p = 0.36). Data are from n = 8 mice. Significance levels are indicated in the panels; *p < 0.05, **p < 0.01, ***p < 0.001. Exact statistical values are reported in the Results and Table S1.

Because the early synchronous Up state coincided with the stimulation-evoked gamma burst described above, we next asked whether this period was associated with enhanced inter-regional coupling. Wavelet coherence revealed a short-latency increase in fast-frequency coupling between PFC, RSC and SUB within the early post-stimulation window (20-100 Hz, 10-100 ms; Fig. 4C). Quantification confirmed that fast-frequency coherence was significantly higher in STIM than CTL sessions (two-ways repeated-measures ANOVA, main effect of Condition: F_(1,14)_ = 7.74, p = 0.03, ηp² = 0.53). Subsequent analyses therefore focused on this early gamma-band window, corresponding to the transient coordinated Up state induced by CLAsc activation.

To determine whether this increase in fast-frequency coherence reflected coordinated transient events rather than a non-specific broadband change, we quantified gamma-burst co-occurrence across regions before and after stimulation. Near-synchronous gamma-bursts, defined as events occurring with less than 10 ms lag and less than 2 Hz frequency difference, occurred more frequently after CLAsc activation across all region pairs (Fig. 4D; paired tests with Benjamini-Hochberg correction, q < 0.05). Thus, the coherence increase corresponds to the alignment of discrete gamma transients across both cortico-cortical and cortico-subicular pairs.

We then quantified cross-regional gamma phase alignment using the Kuramoto index across PFC, RSC and SUB. CLAsc stimulation produced a sharp, time-locked increase in cross-regional gamma phase alignment, peaking approximately 30 to 50 ms after stimulation onset and returning to baseline within the first 100 ms (cluster permutation, p = 0.0075 and p = 0.02; Fig. 4E, Fig. S8A, Table S1). Together with the co-occurrence analysis, this indicates that CLAsc activation does not simply increase local gamma-band activity, but transiently aligns distributed cortico-subicular sites within a common temporal frame.

Finally, we tested whether this brief gamma-coordinated state was accompanied by directed interactions across regions. We computed single-trial covariance-based Granger causality ^25^ on stimulation-aligned gamma signals from PFC, RSC and SUB. CLAsc activation induced a short-latency increase in lagged directed statistical dependence within the first approximately 100 ms, with significant effects from RSC to PFC and from PFC to SUB relative to control sessions (cluster permutation; RSC-to-PFC: p = 0.034; PFC-to-SUB: p = 0.047, p = 0.047 and p = 0.047; Fig. 4F, Table S1). A significant decrease in lagged directed dependence from SUB to RSC emerged over a later time window (cluster permutation, p = 0.03; Fig. 4F, Table S1). By contrast, the instantaneous component, although dynamically modulated around stimulation, did not differ significantly between conditions for any regional pair (Fig. S8B). These results indicate that the early gamma-coordinated Up state is accompanied not only by increased synchrony, but also by a transient reorganization of lagged directed interactions across cortico-subicular circuits. To summarize these distributed changes in directional connectivity, we combined the six Granger-causality measures into a single network-state score using principal-component analysis (PC1, 19.6% of variance explained; loadings shown in Fig. S9). For each mouse, PC1 was calculated and oriented using the other seven animals, after which the excluded mouse’s CTL and STIM profiles were projected onto this independently defined axis. CLAsc stimulation increased the resulting PC1 score in all eight mice. An exact within-mouse permutation test, in which CTL and STIM labels were reversed and PC1 orientation was recalculated for each permutation, confirmed a significant positive shift in the network-state score (p = 0.008). Memory performance also improved after stimulation in this group of animals, though this effect was more modest (one-sided one-sample t-test of ΔMI against zero: t_(7)_ = 1.97, p = 0.0445). Seven of the eight mice showed increases in both measures. An exact within-mouse permutation test combining the changes in PC1 score and memory index confirmed a significant joint effect of stimulation (p = 0.008; Fig. 4G). The size of the physiological and behavioural changes did not correlate across animals (ρ = -0.38, p = 0.36), indicating that stimulation reliably shifted both measures in the same direction without producing a graded, animal-by-animal relationship between them.

Together, these findings show that CLAsc activation recruits a distributed cortical-subicular motif during SWS: a brief synchronous gamma-rich Up state, associated with increased coherence, gamma-burst co-occurrence, phase alignment and directed cortico-subicular interactions, followed by a coordinated Down state, and accompanied by a concordant improvement in memory performance. This sequence provides a candidate physiological mechanism through which CLAsc activity may transiently organize cortical dynamics and hippocampal output during sleep.

## Discussion

We identify a projection-defined claustral population (CLAsc) that links distributed cortical regions with a major hippocampal output node during sleep. CLAsc neurons were preferentially recruited during SWS, and their endogenous activity was organized across coordinated cortical-subicular Up and Down states. Activating this population during post-learning SWS enhanced spatial memory consolidation and imposed a stereotyped network motif consisting of a brief gamma-rich Up state followed by a coordinated Down state across prefrontal, retrosplenial, and subicular regions. During the gamma-rich phase, coherence, gamma-burst synchronization, phase alignment, and lagged directed interactions were transiently reorganized. These findings extend the role of the claustrum beyond cortical-state regulation and identify a pathway through which claustral activity can coordinate cortical dynamics with hippocampal output during sleep.

The claustrum has long been proposed to coordinate distributed cortical activity through its dense and widespread connectivity with the cortical mantle ^1–4^. Previous sleep studies have established the claustrum as a regulator of cortical dynamics, with claustral activity increasing during synchronized states and a retrosplenial-projecting subpopulation modulating cortical delta-wave timing and facilitating memory consolidation ^6,8,9^. Our findings extend this framework by identifying a route through which the claustrum can engage the hippocampal system during sleep. Although the claustrum does not project to the hippocampus proper ^14^, its innervation of the subicular complex provides direct access to a major hippocampal output node ^17,26^. The claustrum may therefore influence hippocampo-cortical communication by coordinating distributed cortical networks with hippocampal output.

Because CLAsc neurons collateralize broadly across cortical territories, including retrosplenial cortex, the population identified here may partially overlap with the retrosplenial-projecting neurons examined in our previous work ^9^. Such overlap would suggest that claustral control of cortical dynamics and hippocampal output may be implemented by partially shared ensembles rather than fully segregated pathways, although direct intersectional mapping will be required to test this possibility.

CLAsc activation improved spatial memory without detectable changes in ripple occurrence or slow oscillation-spindle coupling, indicating that the behavioral benefit is not mediated through coarse modulation of the canonical slow oscillation-spindle-ripple sequence ^27–34^. Our photometry data suggest instead that CLAsc activity is embedded in the ongoing microstructure of SWS. CLAsc activity increased across Up states and around subicular ripples and retrosplenial spindles, and decreased across Down states and following retrosplenial delta waves. These relationships indicate that CLAsc neurons are recruited within naturally occurring SWS network configurations, rather than acting as an autonomous event generator. However, this recruitment is unlikely to reflect a simple linear drive from cortex to claustral output neurons: recent work shows that ACC inputs predominantly suppress claustral regular-spiking neurons through PV-mediated feedforward inhibition ^35^. Thus, endogenous CLAsc recruitment may depend on the convergence of cortical and hippocampal-output signals, together with state-dependent gating within claustral circuits. Such recruitment may also involve changes in the coordination of claustral firing that cannot be resolved by fiber photometry, as recently reported for a distinct claustro-cortical loop during waking behavior ^36^. In turn, our optogenetic data show that activating this population is sufficient to impose a fast gamma-linked Up-to-Down motif across cortico-subicular circuits. CLAsc may therefore operate as a state-dependent feedback node, recruited within specific SWS configurations and capable of transforming this recruitment into a coordinated cortical-subicular state transition.

A key feature of the CLAsc-evoked cortical-subicular motif is the brief distributed gamma burst that precedes the coordinated Down state. Claustral activation has previously been shown to suppress cortical activity through recruitment of inhibitory interneurons ^5,6,11,37^. In this context, the brief gamma-band transient observed here may reflect the rapid engagement of inhibitory network dynamics across cortical and subicular circuits ^38–40^, with the subsequent Down state reflecting its slower population-level consequence. The emergence of interregional synchronization tens of milliseconds after the stimulation pulse is consistent with such local circuit processing and argues against a purely instantaneous stimulation artifact. This interpretation is also consistent with recent human intracranial evidence that slow oscillations can organize gamma-band coordination during memory processing ^41^.

This gamma-coordinated window was accompanied by a transient reorganization of lagged directed interactions. Sleep-dependent hippocampo-cortical communication is most often framed as hippocampal output driving cortical targets, particularly during ripple associated hippocampo-prefrontal coupling ^42,43^. Our Granger causality analysis reveals a complementary and temporally restricted mode: within the first approximately 100 ms after CLAsc activation, lagged directed interactions increased from retrosplenial cortex to prefrontal cortex and from prefrontal cortex to subiculum. The selectivity of this effect to the lagged rather than the instantaneous component argues against a purely zero-lag explanation such as volume conduction. However, the lagged effects may reflect a combination of interregional interactions and region-specific transformations of a common claustral input. Although Granger causality establishes directed statistical dependence rather than direct synaptic causation, this cortico-subicular sequence is consistent with an emerging view that cortical networks can exert temporally structured influence over hippocampal dynamics during sleep ^44–46^, extending it to hippocampal output circuitry. This mode of interaction has remained largely unexplored during sleep and may complement the hippocampus-to-cortex transfer emphasized in current consolidation models. Consistent with this, the distributed reconfiguration of directional connectivity accompanied the memory-enhancing effect of CLAsc stimulation at the group level, although the magnitude of the two effects did not correlate across animals. This dissociation raises the possibility that this network state acts as a permissive substrate supporting consolidation, rather than a graded, quantitative determinant of subsequent memory performance.

More broadly, our findings support an emerging view in which projection-defined claustral populations organize state transitions across distinct networks and behavioral contexts. ACC–projecting claustral neurons have recently been shown to regulate local cortical–state transitions during waking behavior ^36^, whereas our results demonstrate a similar role for CLAsc population in coordinating distributed cortical and subicular dynamics during sleep. Although the underlying circuit mechanisms may differ, these observations suggest that state-transition control may constitute a general computational function of projection-defined claustral pathways.

Several limitations should be noted. Our experiments establish sufficiency rather than necessity: CLAsc activation during post-learning SWS is sufficient to improve memory and evoke the gamma-rich Up-to-Down motif, but loss-of-function experiments are required to determine whether endogenous CLAsc activity is necessary for normal consolidation. Our stimulation was also delivered in open loop within SWS. This establishes that CLAsc activation can bias sleep network dynamics when delivered during SWS, but does not determine whether its efficacy is further gated by the timing of endogenous events such as slow-oscillation phase, spindles, ripples, or Up and Down state transitions. Defining such temporal windows will require future closed-loop experiments. Finally, our recordings were restricted to PFC, RSC and SUB. The subiculum was chosen because it is the principal hippocampal output node targeted by CLAsc neurons, but these recordings do not resolve how the motif relates to activity within hippocampal subfields such as CA1, CA3 or dentate gyrus. Future recordings combining subicular and intrahippocampal sites will be needed to define how CLAsc-dependent cortico-subicular coordination interacts with hippocampal circuit dynamics.

Together, our findings support a model in which the claustrum links cortical and hippocampal-output circuits during slow-wave sleep through a gamma-associated state transition embedded within the Up-Down cycle. CLAsc neurons are recruited within coordinated SWS configurations and, when activated, impose a brief gamma-rich Up state in which cortical-subicular interactions are transiently reorganized, followed by a coordinated Down state. This network motif accompanies enhanced memory consolidation and provides a candidate mechanism through which claustral recruitment may shape distributed sleep dynamics in support of memory.

## Methods

### Animals

All experimental procedures were approved by the local ethical committee for animal experimentation (CREMEAS) and the French Ministry of Research (APAFIS#20388-2019042517013497 and #47167-2024013112207750). A mapping cohort was used for histology only (3 CD1 mice, 1 male, 2 females). Fiber-photometry (claustral calcium imaging) experiments were performed in 7 CD1 mice (4 males, 3 females). Behavioral optogenetic experiments were conducted in 10 CD1 mice (4 males, 6 females), and electrophysiological recordings were obtained from 8 of these animals.

All mice were 8-12 weeks old at the start of the experiments. Environmental conditions were maintained at 24 ±1°C, 30 ±5% humidity, and a 12 h:12 h light/dark cycle (lights on at 08:00), with ad libitum access to food and water. Background noise (45 ±5 dB) was continuously present.

### Surgeries

Mice were anesthetized with isoflurane (4% for induction, 1-2% for maintenance) and placed in a stereotaxic frame. Core temperature was maintained at 37°C using a feedback-controlled heating pad. Local anesthesia (Lurocaïne + Bupivacaïne, 2 mg/kg, s.c.) and systemic analgesia (Metacam, 10 mg/kg, s.c.) were administered before incision. The scalp was disinfected with 70% ethanol and Betadine, the skull was exposed, and the head adjusted to a flat-skull position. In the following section, the stereotaxic coordinates for antero-posterior (AP) and medio-lateral (ML) axes are determined from Bregma, and the dorso-ventral (DV) coordinates are determined from Dura.

#### Viral injections

To selectively target claustral neurons projecting to the subicular complex, all mice received a retrograde viral injection (pAAV.Ef1a-mCherry.IRES.Cre, Addgene #55632-AAVrg; gift from Karl Deisseroth; 0.4 µL per site) targeting the CA1-subiculum region of the hippocampal formation (AP -2.46 mm, ML ±1.75 mm, DV -1.6 mm). For clarity, these neurons are hereafter referred to as subicular-projecting claustral neurons (CLAsc). Secondly, Cre-dependent AAVs were injected into the claustrum (AP +1.25 mm, ML ±2.45 mm, DV -2.50 mm; 0.2 µL per site). The projection mapping cohort was injected only in the right hemisphere and received an AAV5 virus expressing eYFP (pAAV-Ef1a-DIO-eYFP, Addgene #27056-AAV5; gift from Karl Deisseroth). The optogenetic cohort was bilaterally injected with AAV5 virus expressing the blue-light-sensitive excitatory opsin ChETA fused to eYFP (pAAV-Ef1a-DIO-ChETA-eYFP, Addgene #26968-AAV5; gift from Karl Deisseroth, ^47^). The photometry cohort received an AAV9 expressing GCaMP6f (Addgene #100833-AAV9; gift from Douglas Kim and the GENIE Project, ^48^). The syringe was left in place for 10 min after each injection to minimize backflow.

#### Optical element implantation

For the optogenetic cohort, custom-made optical implants were built from 200 µm core-diameter fibers (Thorlabs #FT200EMT) inserted into ceramic ferrules (Thorlabs #CFLC230-10) and positioned bilaterally above the claustrum (AP +1.25 mm, ML ±2.45 mm, DV -2.25 mm). Light transmission efficiency was measured before implantation, and laser output was adjusted to deliver 5 mW at the fiber tip. As for the photometry cohort, a single optical cannula (400 µm core, RWD #R-FOC-BL400C-50NA) was implanted above the right claustrum (AP +1.25 mm, ML ±2.45 mm, DV -2.80 mm). Implants were fixed with dental resin and cement.

#### Electrodes implantation

Both the optogenetic and photometry cohorts received identical LFP electrode implantations in the right hemisphere. Custom platinum-iridium wire electrodes (PHYMEP #778000) targeted the prefrontal cortex (AP +1.70 mm, ML -0.30 mm, DV -1.71 mm), retrosplenial cortex (AP -1.46 mm, ML -0.63 mm, DV -0.89 mm), and subiculum (AP -3.52 mm, ML -2.10 mm, DV -1.23 mm). A cortical ECoG screw was placed over the left hemisphere, with additional screws over the left and right occipital cortices serving as reference and ground. All electrodes were connected to a 16-channel Electrode Interface Board (EIB-16, Neuralynx #31-0603-0106).

### Histology and tissue processing

At the end of experiments, mice were deeply anesthetized (Xylazine 30 mg/kg and Ketamine 200 mg/kg, i.p.) and perfused transcardially (5 min: PBS 0.1 M containing Heparin, pH 7.4; 10 min: Paraformaldehyde-PFA 4%). Brains were post-fixed in PFA 4% overnight at 4°C, then transferred to 20% sucrose in PBS for 48 h at 4°C. Tissue was briefly frozen in isopentane (−40°C, 1 min) and stored at -80°C. Coronal sections (30 µm) were cut at -20°C (NX70, Epredia CryoStar) and mounted to preserve the rostrocaudal extent from prefrontal to entorhinal cortex. Slides were rinsed in PBS (3 × 10 min), blocked for 1 h in 5% horse serum, and incubated overnight at room temperature with gentle agitation (250 rpm) in PBS +2% horse serum containing anti-GFP polyclonal antibody (Alexa Fluor 488; Invitrogen #A-21311, 1:200) and, when applicable, anti-TLE4 antibody (Alexa Fluor 647; Santa Cruz #sc-365406, 1:500). After rinses, sections were counterstained with DAPI and coverslipped with Fluoromount-G (Southern Biotech). Images were acquired on a NanoZoomer (Hamamatsu) and a Zeiss Axio Imager M2 (HXP 120V). QuPath (v0.5.1) was used to assess viral expression and implant locations for all experimental mice.

For claustral projection quantification (anatomical cohort), images were exported as uncompressed OME-TIFF files, registered to the Allen Mouse Brain Atlas using ABBA v0.10.4 with affine and non-rigid spline transformations, and analyzed in Fiji (Bio-Map/Atlas plugin) and QuPath to extract mean GFP fluorescence in each anatomically defined region. Ipsilateral fluorescence was quantified as the mean GFP fluorescence weighted by the number of annotated pixels/objects contributing to each Allen region, and expressed as a percentage of the total GFP signal measured across all cortical and non-cortical annotated regions in that animal. Cortical areas were then grouped into major anatomical groups. Subicular regions were also grouped into a single subicular complex. Superficial and deep cortical projections were quantified separately by pooling respectively layers 1-4 and layers 5-6. Grouped values are reported as mean ± SEM across animals, with each mouse contributing one value per region or layer bin.

### Object location task

The behavioral task was performed only in the optogenetic cohort and was preceded by habituation to the experimenter for at least two weeks. On each testing day, animals were acclimated to the testing room for at least 15 min before the start of the procedure. The open-field arena (square or hexagonal, counterbalanced across within-subject conditions) contained a fixed spatial cue on one wall and was illuminated by indirect LED lighting (30 lux at the center). Continuous background noise was provided by a radio (45 dB at the center). Mice underwent a 10 min habituation session once daily for three consecutive days, with an object placed in the center of the arena on days 2 and 3. During the sampling phase, animals explored two identical objects for 4 min. The recall test occurred 24 h later, during which the same objects were presented for 10 min, but the object least explored during acquisition was displaced to an adjacent corner. Between trials, the walls, floor, and objects were cleaned with 35% ethanol. Memory performance was quantified by measuring the exploration time of each object during test period. The mice were considered to be exploring an object whenever their head was oriented toward and located within 2 cm of the objects. Exploration time was measured manually by the same experimenter from video files. This protocol is designed with an intended time-limited sampling to foil recall of the spatial configuration of the objects on the next day during test period (experimental design inspired from ^42^).

During the 2-3 h post-sampling period, mice remained in the experimental room for a sleeping period where they received light stimulations for claustral activation during the first hour of slow-wave sleep (SWS). Each mouse completed both conditions, STIM (blue-light activation) and CTL (red-light control), in a counterbalanced within-subject design. Mice were connected to electrophysiological device at each step of the behavioral procedure but only the electrophysiological signals recorded throughout the post-sampling sleeping period were used.

### Signal recording

#### Electrophysiological recording

Mice were equipped with a 16-channel amplifier headstage with an integrated three-axis accelerometer (RHD2216, Intan Technologies #C3335) and connected to an Intan Recording Controller (512-channel, #3004, Intan Technologies) operated through the RHX Data Acquisition software. Signals were acquired at 20 kHz and band-pass filtered online between 0.1 and 10 kHz.

#### Optogenetic stimulations and recording

After the sampling phase of the object location task, continuous LFP, ECoG, and accelerometer signals were used for online visual vigilance-state detection. Optogenetic stimulation was manually triggered at the onset of each SWS episode and paused whenever the animal awoke or entered REM sleep, until a total of 60 minutes of SWS stimulation had been reached. Blue light (STIM condition; 473 nm) or red light (CTL condition; 652 nm) was applied in 5 ms pulses at 0.5 Hz. Stimulation timing and duration were recorded via the Intan controller (512-channel, #3004) for precise alignment with electrophysiological data.

#### Fiber photometry recording

Simultaneous recording of fiber photometry and electrophysiological signals were performed for two sessions of 2 to 3 h in their home cage. Dual-color photometry was performed using a custom single-fiber setup equipped with two excitation LEDs: a 470 nm source (M470L4, Thorlabs) to excite GCaMP6f and a 405 nm source (M405LP1, Thorlabs) for the isosbestic control channel. Each LED was modulated by an independent TTL square wave at distinct frequencies (470 Hz and 170 Hz, respectively) generated by a function generator (Rigol DG822), allowing subsequent demodulation of the two signals. Excitation light was collimated with an aspheric condenser lens (ACL25416U-A, Thorlabs), filtered (F47-470, AHF; MF542-20, Thorlabs), and combined using dichroic mirrors (MD453, Thorlabs). A dual-line dichroic beamsplitter (F58-486, AHF) coupled the excitation into the objective (F240FC-A, Thorlabs), while fluorescence emission was directed through the same optical path and detected by a photodiode (DFD_FOA_FC, Doric).

Excitation power at the fiber tip was calibrated between 20 and 100 µW. The photometry signal was digitized through the same Intan Recording Controller (512-channel, #3004, Intan Technologies) used for electrophysiology, ensuring sub-millisecond temporal alignment between fluorescence and LFP recordings.

### Analysis

All analyses were performed in MATLAB using custom scripts adapted to the present datasets.

#### Electrophysiological preprocessing

Raw LFP and ECoG traces were converted to millivolts and downsampled to 2 kHz. Transient artifacts were removed according to the procedure described by ^49^, and line noise (50 Hz and harmonics) was suppressed with adaptive multitaper regression (Chronux ‘rmlinesc.m’ ^50^). For each region, a single representative channel was selected: for PFC and SUB, the least noisy channel, and for RSC, a superficial channel to minimize contamination from the corpus callosum. To extract oscillatory components, signals were decomposed using ensemble empirical mode decomposition (EEMD ^21^; with a noise standard deviation of 0.5 and 20 realizations). From the resulting intrinsic mode functions (IMFs), the fastest (noise) and several slowest (drift) components were discarded. The sum of the remaining IMFs yielded a cleaned broadband signal used for further analyses. Vigilance states were determined manually using combined LFP patterns and accelerometer data and segmented with the *csc-eeg-tools* package (https://github.com/CSC-UW/csc-eeg-tools). Multi-unit activity (MUA) was extracted by filtering the raw signal between 500 and 8000 Hz using a 120^th^-order finite-impulse-response (FIR1) filter. The envelope of the filtered trace was resampled at 2 kHz to match the LFP sampling rate. For optogenetic cohort, only concatenated SWS episodes were analyzed.

#### Photometry preprocessing

The 470 nm and 405 nm channels were demodulated using quadratic demodulation and low-pass filtered (fourth-order Butterworth filter). Signals were then downsampled to 2 kHz, and the initial 3.5 s of each recording were discarded empirically. Photobleaching was corrected by fitting and subtracting a mono-exponential decay function. For each session, both channels were normalized using a z-score based on their combined mean and standard deviation, providing a common reference for comparison across animals and vigilance states.

#### Claustral activity across vigilance states

To quantify claustral calcium activity across vigilance states (Wake, SWS, REM, and Transitions between SWS and REM), the area under the curve (AUC) of both GCaMP6f and isosbestic signals was computed for each concatenated vigilance state. For each mouse, results from the two recording sessions were averaged to obtain a single value per state.

#### Power spectrum slope and theta-delta ratio computation

The power spectrum slope was computed following ^51^. LFPs were divided into 1 s windows with 0.5 s overlap and tapered with a Hamming window. The power spectral density (PSD) of each segment was estimated using Welch’s method, converted to log-log coordinates, and fit with a linear regression between 4 and 100 Hz. The negative of the regression coefficient was taken as the PSS, such that higher values reflect a more marked low frequency dominance. As for Theta-Delta ratio, spectrogram of LFPs were created based on 2 s windows with a step of 2s and a ratio was calculated based on the sum of theta power comprised between 6 Hz and 12 Hz divided by the sum of delta power comprised between the 0.5 Hz and 4 Hz. ***Event analysis.*** All events (delta waves, ripples and spindles) were detected during SWS specifically. *Delta waves* were detected following the criteria of ^52^based on event duration, amplitude, waveform shape, and decrease in fast activity. Briefly, delta wave candidates were determined on the filtered signal (slow signal) between 0.5 and 15 Hz (FIR filter with an order of 1024) by selecting positive waves lasting 0.15 to 0.45 s, with an amplitude exceeding 2 standard deviation (SD) of the z-scored slow signal, and associated with a decrease in fast oscillatory activity (signal filtered between 15 and 200 Hz using a FIR filter and an order of 400). Final delta waves were selected by using a classifier trained to identify delta-wave with a correct waveform shape based on visual validation.

*Spindle* detection was based on the methods described by ^53^ using event amplitude, cycle numbers, and delay between two events. Briefly, the signal was filtered between 9 and 16 Hz using a 600^th^-order FIR filter and squared to obtain power. Spindle candidates were selected if 3 consecutive cycles exceeded 2 SD of the signal power. Each spindle candidate was extended by one cycle at its beginning and end. Spindles candidates overlapping or separated by a duration shorter than 10 ms were merged. Starts and ends of the resulting spindles were selected as the first and last value crossing zero.

*Ripples* were detected following the methods described in ^42,54^, using the FMA toolbox (https://fmatoolbox.sourceforge.net/). Briefly, ripples were isolated from the normalized squared signal filtered between 100 and 250 Hz (FIR filter, with an order of 60) by selecting bursts of activity exceeding 2 SD but below 5 SD. Bursts separated by less than 30 ms were merged, and the resulting ripples-candidates with a duration lower than 20 ms or higher than 100 ms were discarded. Finally, for each animal, contralateral ECoG was used as a noise control.

*Claustral response to events -* Photometry response to single event was calculated by z-scoring claustral activity within a 4 s window around each event center and averaging them across recording sessions and mice. The displayed signal represents the average signal across all sessions for each mouse.

*Event probability -* To compute peri-stimulus event probability, for each time point around a given repeated trigger (light stimulation or single event center), the presence (1) or absence (0) of event was summed and divided by the total number of trigger repetitions.

#### Co-distribution analysis

Signal X values (PSS, theta-delta ratio, event rate) from each region were divided into deciles, and the mean fluorescence associated with each decile was computed. Variability was estimated using bootstrap resampling (500 iterations). Mean fluorescence values were then averaged across bootstrap iterations to obtain robust estimates of fluorescence variations across the range of signal X values. Analyses were performed separately for GCaMP and isosbestic signals.

#### Electrophysiological response to claustral stimulation

Mean LFP responses were obtained by aligning traces to stimulation onset and averaging across all pulses.

#### Stimulation-locked phase consistency within regions

Phase consistency across stimulations was quantified using the Kuramoto index ^55,56^. Instantaneous phase was extracted from the Hilbert transform of each trial’s analytic signal. The Kuramoto index K(t) was computed across trials at each time point to quantify trial-to-trial phase alignment. The Hilbert transform was applied to each LFP trace, segmented between -0.2 s and +0.6 s around stimulation onset. The Kuramoto index *K*(*t*) was computed as

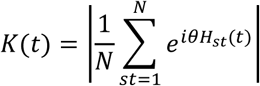

Where *N* is the total number of stimulations and *θH_st_(t)* is the instantaneous phase of the Hilbert transformed signal for each stimulation *st*. Values range from 0 (no phase consistency) to 1 (perfect synchrony). If the claustrum stimulation induces a consistent response in cortical regions, the different traces should have a similar phase and *K* should increase.

#### Time-frequency analysis

Frequency-dependent amplitude changes were calculated after time-frequency transformation using the continuous wavelet transform implemented in Matlab (cwt). The transform was computed on the LFPs data using the sampling frequency and restricting the analysis to frequencies between 0.1 and 300 Hz. (morse wavelet, 10 voices per octave).

#### Gamma-composite

Composite gamma signals were reconstructed by summing the intrinsic mode functions (IMFs) whose peak frequencies lay between 25 and 200 Hz, providing a broadband but physiologically constrained gamma trace.

#### Gamma-burst detection

Gamma bursts were detected using an adaptation of the approach of ^23^, modified to operate on the entire concatenated SWS recordings rather than on individual theta cycles. Spectrograms of gamma composite were computed with a complex Morlet wavelet transform (0.5 s window, 20-200 Hz in 3 Hz steps, 6-20 cycles). Bursts were extracted directly from the spectrogram as contiguous patches of locally elevated power, identified after grayscale binarization and connected-component labeling ^57^. Detection was repeated across descending power thresholds (100-85% of the maximum) to capture all distinct power peaks while preventing artificial fusions between neighboring events. For each validated component, we computed onset and offset times, duration, median power, and power-weighted frequency and temporal centroids.

#### Up and Down state analyses

Up and Down states were detected using the Explicit-Duration Hidden Markov Model (EDHMM) approach ^24^, applied to the log-transformed MUA envelope and, when available, the high-gamma envelope (>40 Hz). Features were binned into 10 ms non-overlapping windows, averaged within bins, and z-scored across time. Initial clustering was performed by k-means, followed by EDHMM inference enforcing a minimum state duration of 100 ms and alternating transitions. Only states with posterior confidence >90% were retained, and each region’s output was stored as a binary vector labeling Up and Down states.

To measure the Up and Down state modulation around the claustral optogenetic stimulation, normalized probabilities of Up- and Down-states were computed around each stimulation and z-scored within the surrounding window.

Photometry activity around transitions toward Down states was computed within a 2 s window. For events shorter than 2 s, samples outside event boundaries were excluded. Mean fluorescence signals were compared with shuffled transitions. For each region, linear mixed-effects models were used to compare pre-versus post-transition activity between true and shuffled signals. As for photometry activity within Up or Down, episodes shorter than 50 ms were excluded. Remaining episodes were divided into deciles according to their duration, and mean fluorescence was computed for each decile. Fluorescence values were averaged across conditions within each mouse and normalized by subtracting the first decile. For each region, linear mixed-effects models were used to assess the effects of decile, signal type, and state (Up vs Down).

#### Synchronous Up and Down states

Common Up and Down episodes were defined as periods during which all three regions were simultaneously in the same Up or Down state. The resulting synchronous Up and Down states were then considered as individual events and their probability of occurrence was measured around the optogenetic activation, with a z-scored normalization for each window around the stimulation. Photometry distributions during synchronous Up and Down states were computed by binning photometry values (26 bins) and calculating their probability to occur during the common Up or common Down states.

#### Inter-regional coherence

Amplitudes were normalized over time to evaluate frequency-specific modulation. Time-frequency relationships between regions were further assessed by computing magnitude-squared wavelet coherence across all region pairs using the Matlab built-in function (wcoherence; analytic morse wavelet, 0.1300 Hz). Coherence values were normalized over time and averaged across stimulations. Between-condition comparisons were then performed separately for fast (20-100 Hz, 0.01-0.1 s post-stim) and slow (3-7 Hz, 0.1-0.5 s post-stim) frequency bands.

#### Synchronous gamma bursts

For each stimulation, two 50 ms windows (PRE and POST stimulation) were analyzed. A co-occurrence was defined when at least one event per region occurred within a temporal difference (ΔT) and frequency difference (ΔF) satisfying ΔT ∈ {2, 5, 8, 10, 15, 20 ms} and ΔF ∈ {0.5, 1, 2, 5, 10 Hz}. For each region pair, window, and parameter combination, a binary vector was generated (1 = co-occurrence present, 0 = absent). The mean of this vector represented the fraction of stimulations associated with gamma co-occurrence. Group-level values were obtained by averaging across mice.

#### Inter-regional gamma-phase synchronization

The Kuramoto index was computed across regions (PFC, RSC, SUB) at each time point, using the instantaneous phase of the gamma composite. This Kuramoto index was averaged across stimulations to capture cross-region gamma coherence. Because the gamma range overlaps with hippocampal ripple frequencies (∼100 Hz), ripple probability was quantified separately.

#### Covariance-based Granger causality (covGC)

To quantify directed information transfer between brain regions in the gamma frequency band following optogenetic stimulation, we used the method developed by Brovelli and colleagues ^25^, which has been implemented in the open-access FRITES toolbox (https://brainets.github.io/frites/index.html). CovGC evaluates in short time windows whether the past activity of a source region reduces the uncertainty of the current activity of a target region.

The estimation relies on covariance matrices computed from time-lagged observations and assumes that the signals can be approximated by Gaussian processes. Under these assumptions, covariance-based Granger causality provides an information-theoretic estimate of directed interactions that is equivalent to classical Granger causality. Measures of covGC were computed around each optogenetic stimulation using sliding windows of 100 ms with a step size of 10 ms starting 0.5 s before the stimulation and ending 0.8 s after it. Within each window, time-lagged observations were constructed using a total lag of 10 ms, and conditional entropies were estimated from covariance matrices of these lagged signals to quantify directed and instantaneous interactions between regions. The resulting covGC measures were then averaged across stimulations to determine the mean effect of claustral activation on the interregional dependencies.

#### PCA-derived network-state score and relationship to memory performance

To summarize the time-resolved pattern of directed interactions following claustral stimulation, covariance-based Granger-causality responses were extracted for the six directed connections among PFC, RSC, and SUB. For each stimulation trial and directed connection, GC values were normalized relative to their trial-specific pre-stimulation baseline as 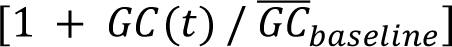. Normalized post-stimulation responses were averaged across trials separately for each mouse and condition, and the six resulting directed GC time courses were concatenated into a single feature vector for each CTL or STIM session. Each directed-connection-by-time feature was standardized across sessions, and principal component analysis was applied to the resulting session-by-feature matrix. The first principal component was retained as a scalar GC-state. For visualization, the full-sample PC1 was oriented such that its mean score was higher in STIM than in CTL sessions, and its temporal and directional loading structure was displayed by reshaping the PC1 coefficients into a directed-connection-by-time matrix. For directional inference, a leave-one-mouse-out procedure was used in which both sessions from one mouse were excluded, feature centering and scaling and PCA fitting were performed using the remaining mice, and PC1 polarity was determined from the mean CTL-to-STIM score difference in this training set; the CTL and STIM profiles of the excluded mouse were then standardized using the training-set parameters and projected onto the independently fitted and oriented PC1 axis, yielding a cross-validated STIM-CTL network-state difference for each mouse.

### Statistical analysis

Statistical analyses were performed using Jamovi (v2.6.26.0) and MATLAB (R2024b), and results are reported in the corresponding figure legends. Assumptions of normality and homoscedasticity for parametric tests were evaluated using the Shapiro-Wilk test and by visual inspection of residual Q-Q and homogeneity plots. When these assumptions were violated, equivalent non-parametric tests were applied. Boxplots display minima and maxima, while time-series data are shown as mean ± SEM. Unless otherwise indicated, the significance threshold was set to α = 0.05. When p-values appear in figures, p < 0.05 (*), p < 0.01 (**), and p < 0.001 (***), denote increasing levels of significance; otherwise, exact p-values are reported. When post-hoc tests were conducted, Bonferroni correction was applied unless otherwise specified.

#### Photometry activity across vigilance states

The AUC of both GCaMP6f and isosbestic channels was analyzed with a linear mixed-effects model including Signal (GCaMP6f vs. isosbestic), State (Wake, SWS, REM, Transition), and Session (two per animal) as fixed factors, and Mouse as a random effect.

#### Co-distribution correlations

To test the statistical relationship between photometry signals and other metrics (PSS, theta-delta ratio, event rate) we fitted for each metric a Generalized Additive Mixed Models (GAMMs), using the *mgcv* package in R (version 1.9-4). Each GAMM included a penalized cubic regression spline (bs = ‘cr’), with a basis dimension (k) fixed to 15 and Signal (GCaMP vs. isosbestic) as a parametric factor. Animal identity was included as a random effect to account for repeated measurements within individuals. The random effect structure (random intercept alone, or intercept + slope over the quantile axis) was selected automatically: a random slope model was first attempted, and retained only if it converged without warnings and yielded a non-negligible slope variance; otherwise, a random intercept model was used. To test whether the smooth relationship with the variable differed between the GCaMP and isosbestic signals, the primary inferential contrast, a likelihood ratio test (LRT) was performed by comparing a full model (signal-specific smooths) to a null model (shared smooth), both fitted by maximum likelihood (ML). Final smooth estimates and standard errors were obtained by refitting the full model using restricted maximum likelihood (REML). The difference between GCaMP and isosbestic smooths was quantified pointwise via posterior simulation (10,000 draws from the multivariate normal posterior of model coefficients), yielding 95% credible intervals on the difference curve.

#### Object-location task performance

Statistical significance for multi-factors comparison (Condition and Object) was assessed using two-way repeated-measures ANOVA followed by Holm-Bonferroni procedure for multiple comparisons. Exploration time and memory index comparisons were measured with a two-tailed paired Student t-test. Comparison to chance level was assessed with a one-tailed one-sample t-test.

#### Paired time-series comparisons

Significance across temporal samples was assessed using cluster-based permutation tests ^58^. A paired t-test was computed at each time point, and clusters of consecutive samples with p < 0.05 and identical sign were identified. Cluster strength was defined as the sum of t-values within each cluster. To obtain corrected p-values, 10,000 random permutations were generated by shuffling condition labels within animals, and the distribution of maximal cluster strengths was used as the null distribution. Observed clusters were considered significant when their strength exceeded the 95^th^ percentile of this null distribution. For photometry activity across Up-to-Down transitions or within the event, linear mixed-effects models were used.

#### Wavelet coherence

Coherence values in the slow (3-7 Hz) and fast (20-100 Hz) bands were analyzed using two-way repeated-measures ANOVAs with Condition (STIM vs. CTL) and Region Pair as within-subject factors.

#### Synchronous Up-Down photometry distribution

Kolmogorov-Smirnov statistics were calculated for GCaMP and isosbestic signals using subsampled Up and Down events and compared with permutation-derived null distributions generated by shuffling pooled samples across conditions. Statistical significance was assessed using two-way ANOVA followed by Bonferroni’s correction for multiple comparisons.

#### Gamma burst co-occurrence

The change in co-occurrence (ΔPOST - PRE) between conditions was tested with paired one-tailed t-tests (H₀: STIM = CTL; H₁: STIM > CTL) across mice, separately for each region combination and each (ΔT, ΔF) parameter pair. The resulting p-values were corrected for multiple comparisons using the Benjamini-Hochberg false discovery rate procedure with a false discovery rate set to q < 0.05.

#### PCA-derived GC-state analysis

The directional effect of stimulation on the cross-validated PC1 score was assessed using an exact fold-wise sign-flip permutation test. All 2^n^ possible reversals of CTL and STIM labels within mice were enumerated; for each permutation, PC1 polarity was re-estimated within each leave-one-mouse-out training set, and the one-sample t-statistic of the reconstructed cross-validated differences was used as the test statistic. To test whether stimulation jointly increased the network-state score and memory performance, the same within-mouse sign flips were applied simultaneously to the physiological and behavioral differences. Separate one-sample t-statistics were computed for both measures and combined as 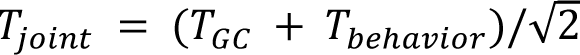 . Exact one-sided p-values were calculated as the proportion of permutations yielding a statistic at least as large as the observed value. The number of mice showing positive changes in both measures was reported descriptively, and the association between the magnitudes of physiological and behavioral changes was assessed using Spearman’s rank correlation.

All data visualization and figure preparation were performed in MATLAB, GraphPad Prism, and finalized in Adobe Illustrator or Affinity Designer for layout and labeling.

## Supporting information

supplementary informations

## Acknowledgements

This work was supported by CNRS, Université de Strasbourg and a grant from the Agence Nationale de la Recherche: ANR CLAwaves (ANR-24-CE37-3417). We acknowledge the In Vitro Imaging Platform – Strasbourg (CNRS UAR3156), member of the national infrastructure France-BioImaging supported by the French National Research Agency (ANR-10-INBS-04). The authors wish to thank Dr. Andrea Brovelli for fruitful discussion around the single-trial Granger causality analysis.

## Author contributions

CP and FT performed the experiments and analyzed the data; MA and TB analyzed the data; KH and CM performed the immunohistochemical analysis; YS supervised the photometry experiments; JJ and DB actively participated in discussions and experimental planning; RG conceived the experimental and analytical design; all authors wrote the article.

