## supplementary informations for "Claustral pathway coordinates distributed cortical dynamics with hippocampal output during sleep to promote memory consolidation"

**<u>Supplementary Information</u>**

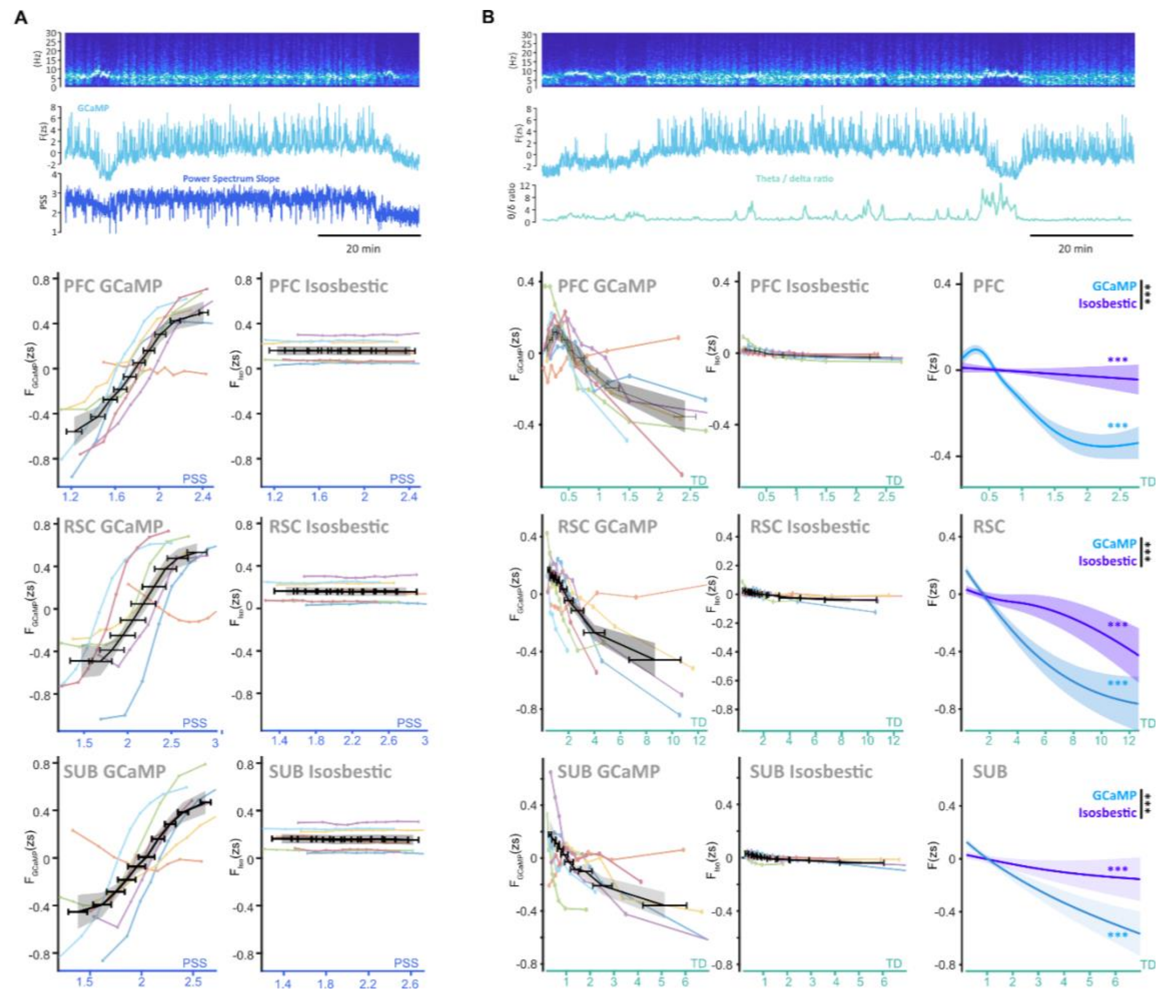


**Figure S1. CLAsc activity covaries with cortical and subicular dynamics.**

**A**. CLAsc GCaMP6f and isosbestic fluorescence as a function of power-spectrum slope in PFC, RSC, and SUB during SWS. Thin colored lines show individual mice (n = 7; values averaged across recording sessions), and black dashed lines show the group mean. GAMM fits comparing the dependence of GCaMP6f and isosbestic signals on power-spectrum slope are shown in the right-hand panels.

**B.** CLAsc GCaMP6f and isosbestic fluorescence as a function of the theta/delta ratio in PFC, RSC, and SUB during SWS. Thin colored lines show individual mice, black dashed lines show the group mean, and the right-hand panels show the corresponding GAMM fits.

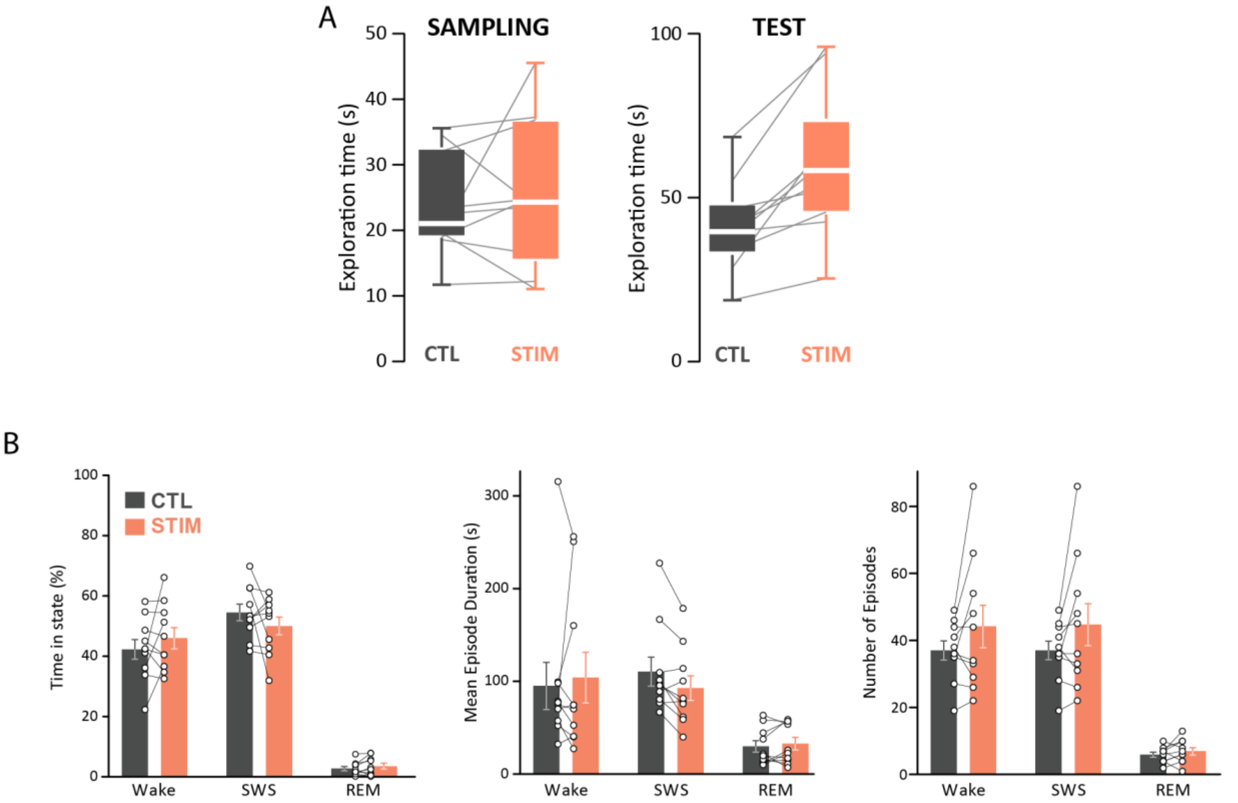


**Figure S2: CLAsc stimulation does not alter object exploration or post-learning sleep architecture.**

**A.** Total object-exploration time during the sampling and test phases under CTL and STIM conditions. Lines connect values obtained from the same mouse.

**B.** Sleep architecture during the first 2 h after learning. From left to right: percentage of time spent in Wake, SWS, and REM sleep; mean episode duration for each vigilance state; and number of episodes. CLAsc stimulation did not significantly alter any of these measures relative to the CTL condition. Bars show mean ± SEM, individual mice are overlaid, and lines connect paired observations.

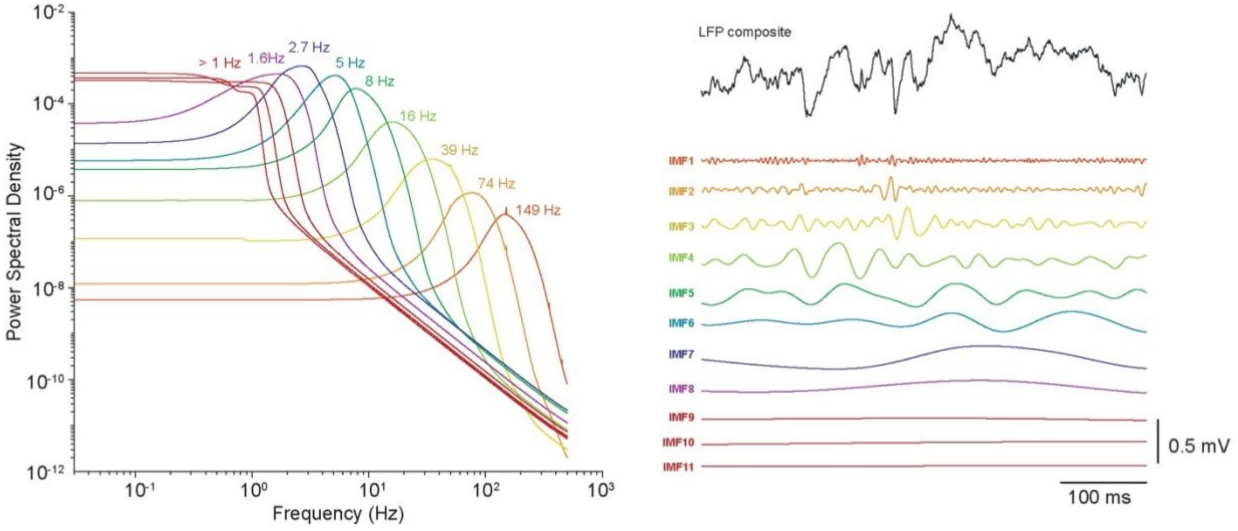


**Figure S3: Signal decomposition using ensemble empirical mode decomposition (EEMD).**

(Left) Power spectra of intrinsic mode functions extracted from representative cortical LFP traces using ensemble empirical mode decomposition. Individual modes span characteristic frequency ranges from slow components below 1 Hz to fast components approaching 150 Hz.

(Right) Representative raw LFP trace and the corresponding intrinsic mode functions, color-coded as in *left.* Ensemble empirical mode decomposition adaptively separates the signal into oscillatory modes without imposing fixed a priori frequency bands, enabling analysis of spontaneous and stimulation-evoked activity across timescales.

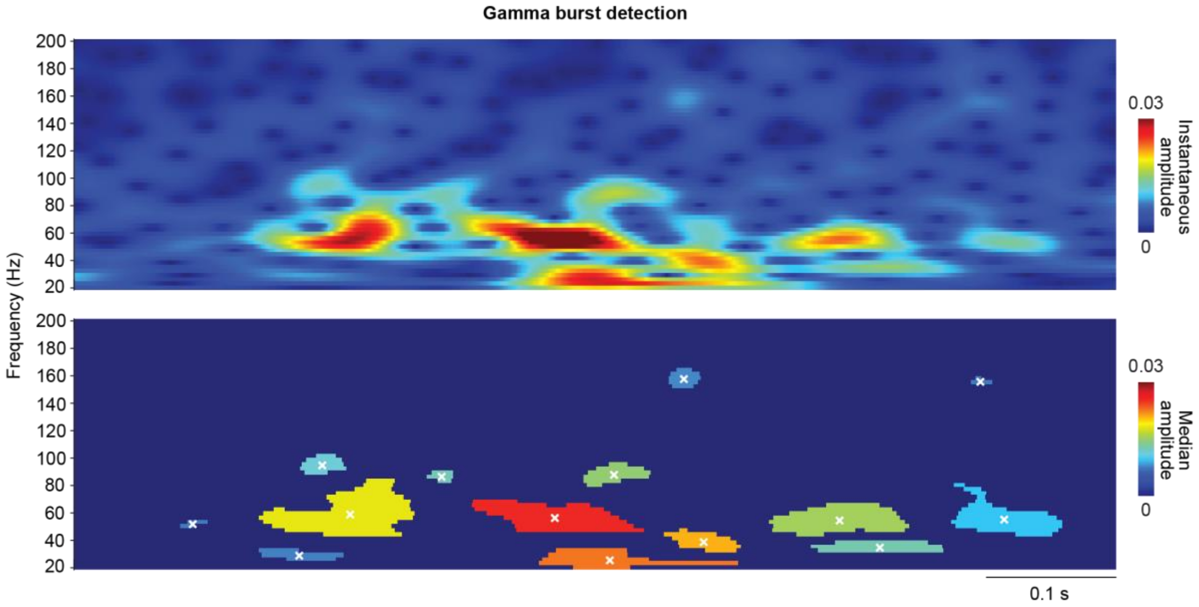


**Figure S4: Detection of gamma bursts in time-frequency space.**

Gamma bursts were detected using an adaptation of the approach described by Douchamps et al. (2024), applied to composite gamma signals reconstructed by summing intrinsic mode functions with peak frequencies between 25 and 200 Hz.

Top. Representative time-frequency map of the composite gamma signal computed using a complex Morlet wavelet transform over the 20–200 Hz range.

Bottom. Corresponding binary map of contiguous power clusters identified after grayscale thresholding and connected-component labeling. Each colored patch represents an individual detected gamma burst. White crosses indicate the power-weighted temporal and frequency centroid used to define the time and frequency of each event.

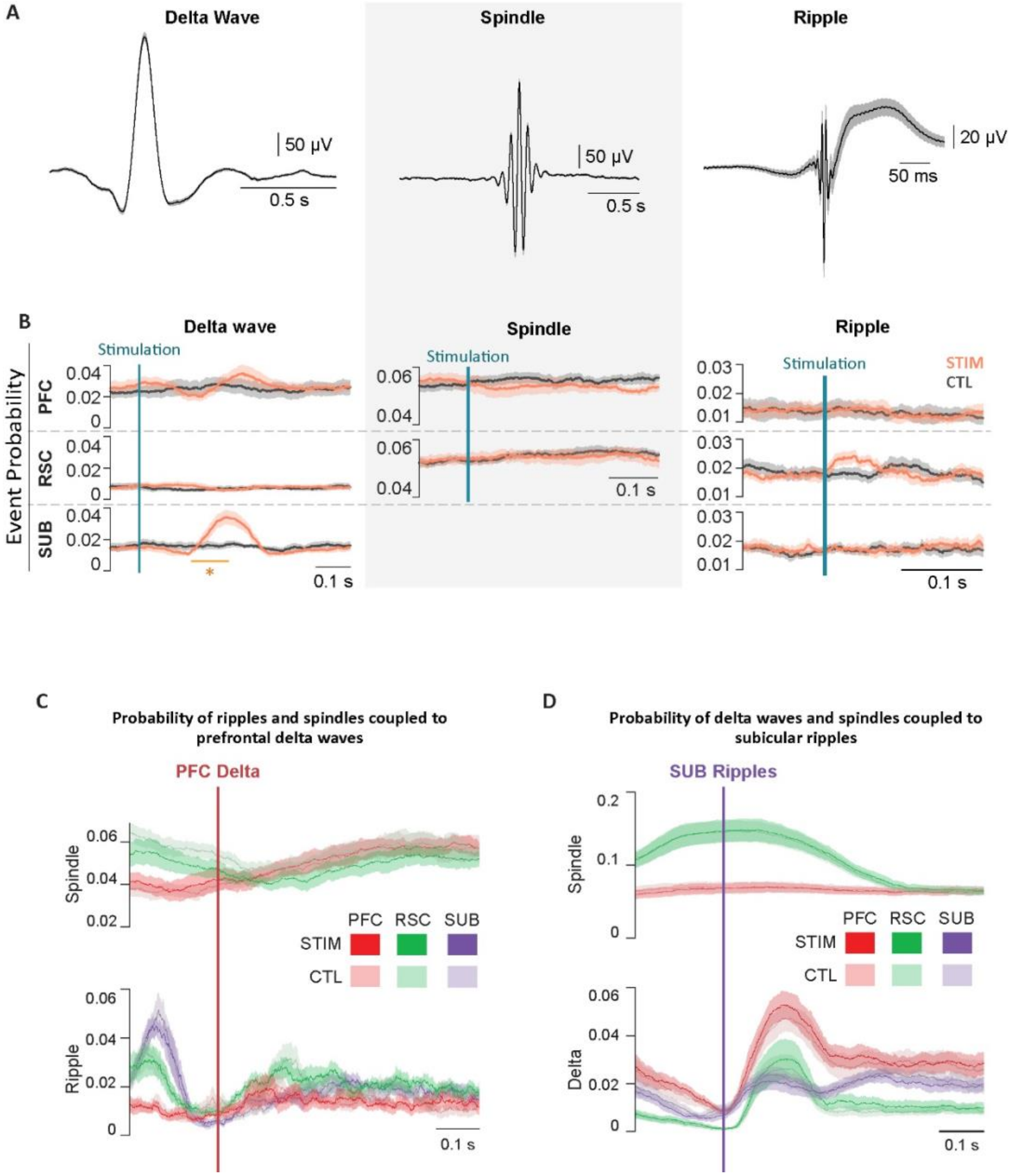


**Figure S5: Probability of delta waves, spindles, and ripples in response to CLAsc stimulation.**

**A.** Mean LFP waveforms aligned to the peak of each detected event type. From left to right: delta waves, spindles, and ripples. Delta waves were detected using criteria based on event duration, amplitude, waveform shape, and reduction in fast activity. Spindles and ripples were detected using the procedures described in the Methods. Delta-wave detection was used to characterize oscillatory coupling and was not used as a proxy for Down states.

**B.** Probability of delta waves, spindles, and ripples aligned to CLAsc stimulation. CLAsc activation did not alter spindle or ripple occurrence but produced a transient increase in delta-wave probability, most prominently in SUB. Significant post-stimulation periods identified by cluster-based permutation tests are indicated by orange bars. Shaded areas show mean ± SEM.

**C.** Event probabilities aligned to prefrontal delta waves under STIM and CTL conditions. No significant differences were observed between conditions. STIM and CTL curves largely overlap because their mean values are similar.

**D.** Event probabilities aligned to subicular ripples under STIM and CTL conditions. No significant differences were observed between conditions. The traces confirm the expected temporal coupling among subicular ripples, cortical spindles, and delta waves. STIM and CTL curves largely overlap because their mean values are similar.

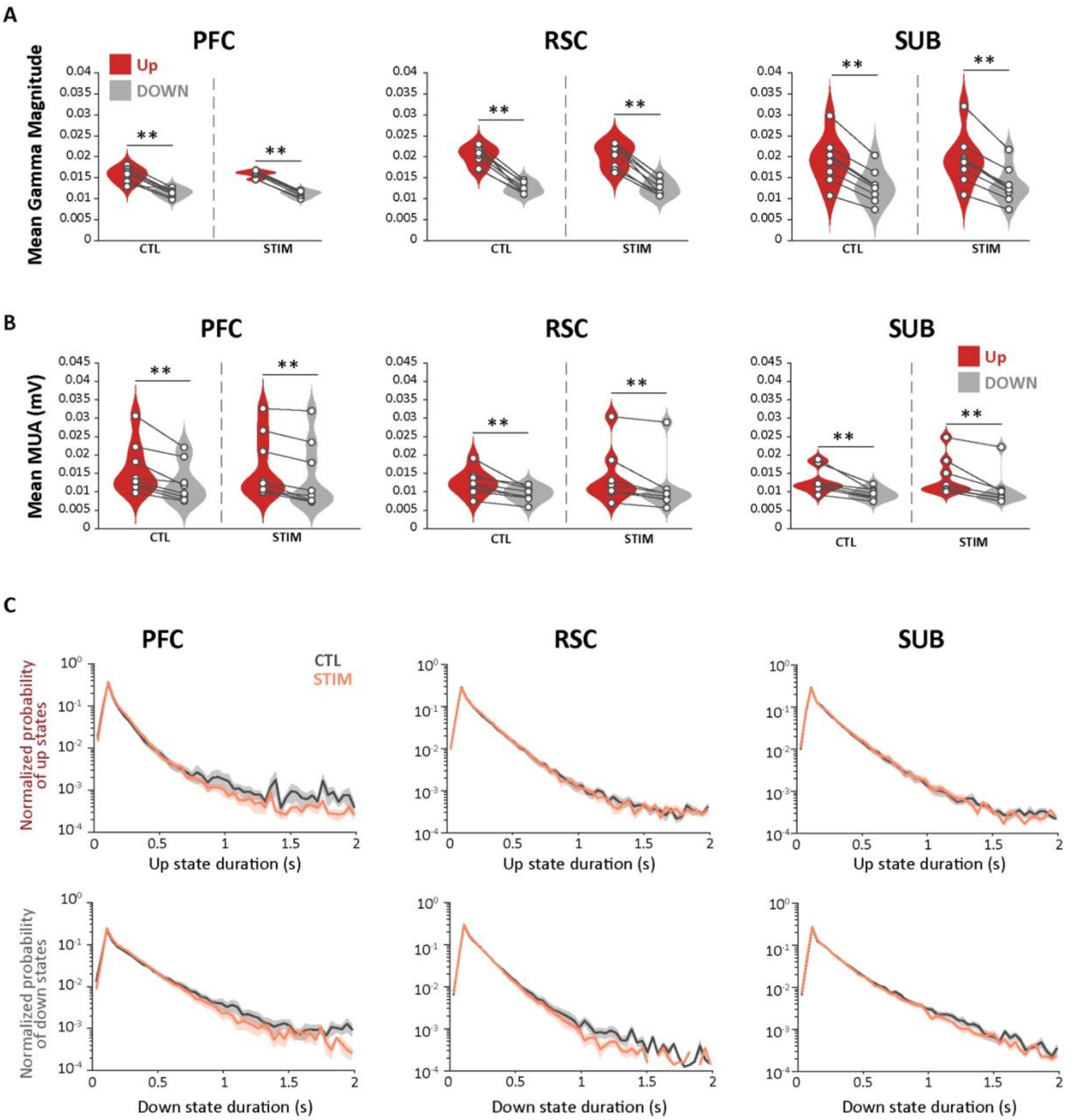


**Figure S6: Validation and duration of Up and Down states in STIM and CTL conditions.**
**A.** Mean gamma magnitude during Up and Down states in PFC, RSC, and SUB under CTL and STIM conditions. Up states showed greater gamma activity than Down states across regions (**$p < 0.01$).
**B.** Mean MUA amplitude during Up and Down states in PFC, RSC, and SUB under CTL and STIM conditions. Up states showed greater MUA than Down states across regions (**$p < 0.01$).
**C.** Probability-density distributions of Up-state and Down-state durations across all recorded sessions under CTL and STIM conditions. Both states displayed characteristic right-skewed duration distributions, with no significant difference in mean duration between conditions.

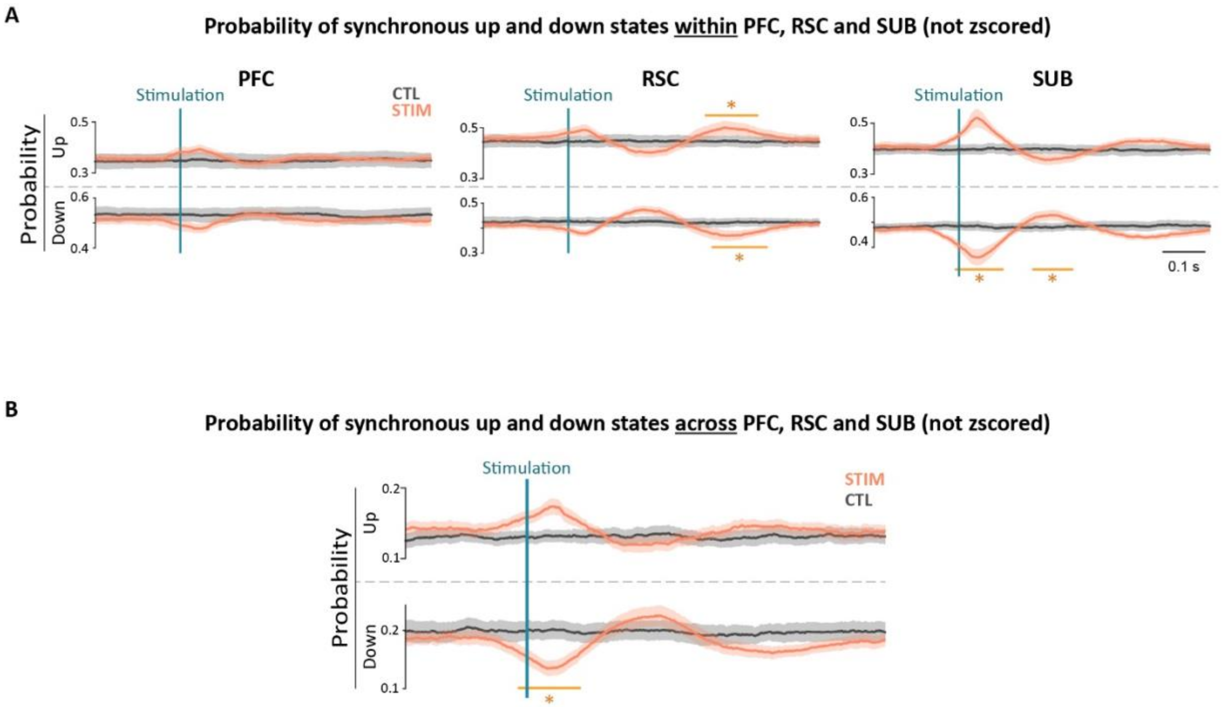


**Figure S7: Non-z-scored probability of local and synchronous Up- and Down-states in response to CLAsc stimulation.**

**A.** Probability of local Up states and Down states aligned to CLAsc optogenetic stimulation in PFC, RSC, and SUB. CLAsc activation transiently increased Up-state probability shortly after stimulation and enhanced Down-state probability approximately 200 ms later, with region-specific significance patterns.

**B.** Probability of synchronous Up states and Down states occurring simultaneously across PFC, RSC, and SUB. A coordinated Up state emerged shortly after CLAsc activation and was followed by a synchronous Down state. Significant post-stimulation periods identified by cluster-based permutation tests are indicated by orange bars. Shaded areas show mean ± SEM.

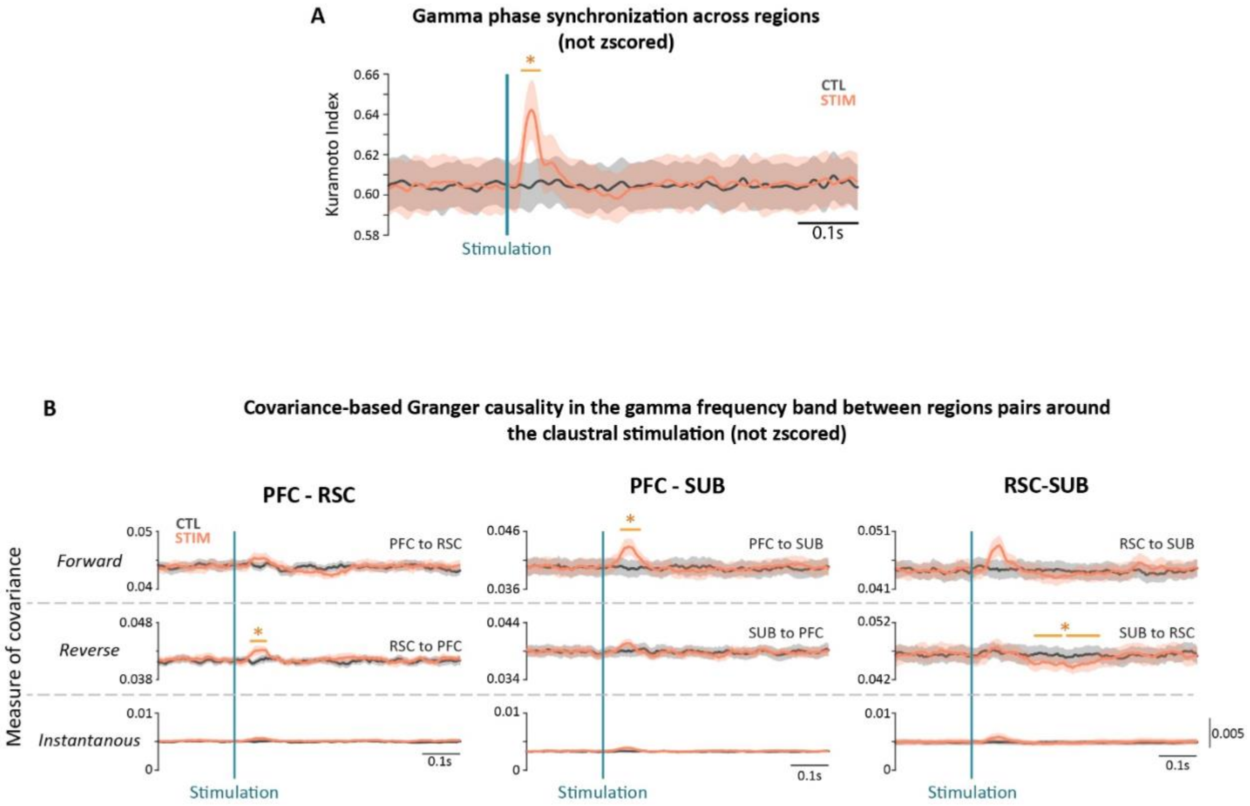


**Figure S8: Non-z-scored gamma-phase synchronization and covariance-based Granger causality following CLAsc stimulation.**

**A.** Mean Kuramoto index computed across PFC, RSC, and SUB in the gamma range and aligned to CLAsc stimulation. A transient increase in cross-regional gamma-phase synchronization occurred shortly after stimulation, consistent with the z-scored analysis shown in Figure 4E. Shaded areas show mean ± SEM across animals.

**B.** Stimulation-aligned covariance-based Granger-causality analysis of gamma-band interactions between PFC, RSC, and SUB. For each region pair, the upper two traces show lagged directed interactions in the forward and reverse directions, whereas the lower trace shows the instantaneous component. CLAsc activation produced short-latency changes in lagged directed interactions, including increases from RSC to PFC and from PFC to SUB. Shaded areas show mean ± SEM; horizontal bars indicate significant STIM versus CTL clusters.

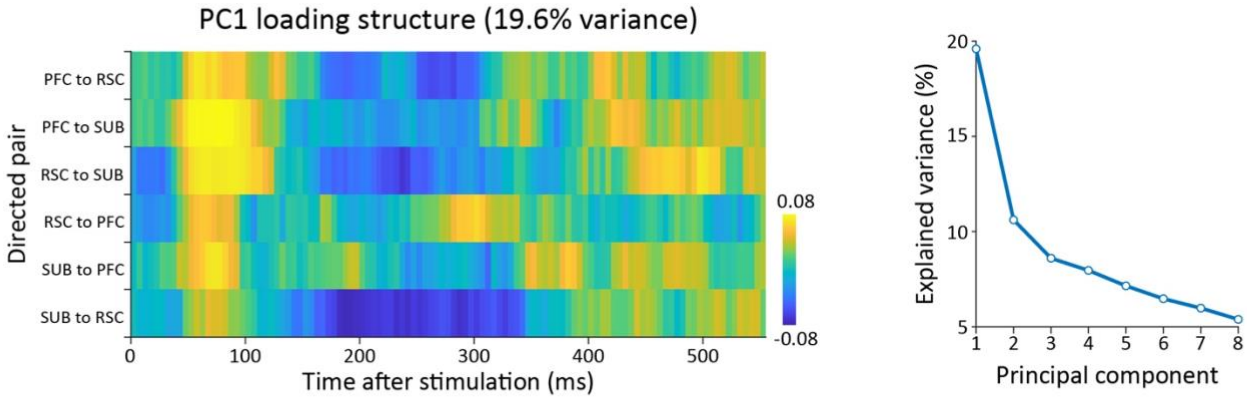


**Figure S9: Principal-component structure of stimulation-evoked Granger-causality changes.**

**A.** Temporal loading structure of the first principal component derived from the six directed Granger-causality measures among PFC, RSC, and SUB. Loadings are shown for each directed connection across the post-stimulation period. The first principal component explained 19.6% of the total variance.

**B.** Scree plot showing the percentage of variance explained by successive principal components**.**

**Table S1. Summary of cluster-based permutation statistics (next page).**

Results of cluster-based permutation tests for all figures. Each row reports the analysis type, brain region (or region pair), cluster mass values, cluster-level p-values, and corresponding effect sizes (Cohen's d). When multiple significant clusters were identified for the same analysis, all are listed within the same row in temporal order (earliest to latest). Positive d values indicate increases in the STIM relative to the CTL condition, and negative values indicate decreases.

| Figure | Analysis | Region | Cluster mass value | Cluster-level p-value | Effect size (Cohen's d) |
|---|---|---|---|---|---|
| **Figure 3A** | LFP response to CLAsc stimulation | PFC | 601.83; 1004 | 0.03; 0.0075 | 1.47; -1.8 |
| | | RSC | 436.86; 630.14; 317.21; 507.36 | 0.03; 0.01; 0.04; 0.01 | 1.27; 1.65; 1.11; -1.15 |
| | | SUB | 12963; 423.27; 373.34; 586.49 | 0.0075; 0.04; 0.04; 0.03 | 1.83; -1.53; -1.33; -1.68 |
| | Kuramoto Index across CLAsc stimulation | PFC | 2395,8 | 0.0075 | 2.03 |
| | | RSC | 516.28 | 0.0075 | 1.03 |
| | | SUB | 3.1891; 405.37 | 0.0075; 0.047 | 1.85; 1.42 |
| **Figure 3B** | Gamma burst after the stimulation | PFC | 31.83 | 0.0075 | 2.13 |
| | | RSC | 48.28; 41.51; 37.61 | 0.02; 0.03; 0.04 | 1.74; 1.38; -1.32 |
| | | SUB | 67.38 | 0.0075 | 2.02 |
| **Figure 3D** | Up-state probability after CLAsc stimulation | PFC | 923.59 | 0.0075 | 2.52 |
| | | RSC | 663.42; 1087.22 | 0.037; 0.01 | 1.13; -1.33 |
| | Down-state probability after CLAsc stimulation | PFC | 836.67 | 0.0075 | -2.23 |
| | | RSC | 1268.23; 707.65 | 0.01; 0.04 | 1.31; -1.18 |
| | | SUB | 1482.79; 597.19 | 0.0075; 0.047 | 1.83; -0.94 |
| **Figure 4A** | Synchronous up-state around CLAsc stimulation | PFC-RSC-SUB | 639.82; 1190.74 | 0.04; 0.01 | 1.34; -1.94 |
| | Synchronous down-state around CLAsc stimulation | PFC-RSC-SUB | 1035.75; 534.71 | 0.014; 0.046 | 1.40; -1.12 |
| **Figure 4E** | Cross-region Kuramoto Index in the gamma band around CLAsc stimulation (zscore) | PFC-RSC-SUB | 284.41; 122.68 | 0.0075; 0.020 | 1.76 2.04 |
| **Figure 4F** | Granger Causality (zscore) | PFC to SUB | 30.10; 30.46; 26.76 | 0.047; 0.047; 0.047 | 1.01; -1.84; -1.67 |

| | | | | | |
|---|---|---|---|---|---|
| | | RSC to PFC | 24.23 | 0.034 | 1.31 |
| | | SUB to RSC | 53.41 | 0.03 | -1.41 |

**Table S2. Summary of cluster-based permutation statistics for supplementary figures.** Same as Table S1 for supplementary figures.

<table>
<tr><th>Figure</th><th>Analysis</th><th>Region</th><th>Cluster mass value</th><th>Cluster-level p-value</th><th>Effect size (Cohen's d)</th></tr>
<tr><td>Figure S5B</td><td>Event response to stim (delta wave)</td><td>SUB</td><td>1170</td><td>0.0075</td><td>1.76</td></tr>
<tr><td rowspan="4">Figure S7</td><td>Up state probability around stim</td><td>RSC</td><td>750.73</td><td>0.049</td><td>1.12</td></tr>
<tr><td rowspan="2">Down state probability</td><td>RSC</td><td>781.00</td><td>0.03</td><td>-1.10</td></tr>
<tr><td>SUB</td><td>637.50;<br>581.07</td><td>0.04;<br>0.04</td><td>1.23;<br>-0.93</td></tr>
<tr><td>Synchronous down-states probability around the stimulation</td><td>PFC-RSC-SUB</td><td>697.08</td><td>0.04</td><td>-1.23</td></tr>
<tr><td>Figure S8A</td><td>Gamma synchronization around stim (Kuramoto index not zscored)</td><td>PFC-RSC-SUB</td><td>271.14</td><td>0.0075</td><td>1.78</td></tr>
<tr><td rowspan="3">Figure S8B</td><td rowspan="3">Granger Causality not zscored</td><td>RSC to PFC</td><td>33.43</td><td>0.03</td><td>1.55</td></tr>
<tr><td>PFC to SUB</td><td>33.75</td><td>0.03</td><td>1.14</td></tr>
<tr><td>SUB to RSC</td><td>58.24;<br>84.51</td><td>0.02;<br>0.0076</td><td>-1.84;<br>-1.83</td></tr>
</table>
